# Structural development of amyloid precursors in insulin B chain and the inhibition effect by fibrinogen

**DOI:** 10.1101/2021.12.26.474222

**Authors:** Naoki Yamamoto, Rintaro Inoue, Yoshiteru Makino, Naoya Shibayama, Akira Naito, Masaaki Sugiyama, Eri Chatani

**Affiliations:** Division of Biophysics, Physiology, School of Medicine, Jichi Medical University, 3311-1 Yakushiji, Shimotsuke, Tochigi, 329-0498, Japan; Institute for Integrated Radiation and Nuclear Science, Kyoto University, 2 Asashiro-Nishi, Kumatori, Sennan-gun, Osaka, 590-0494, Japan; Graduate School of Engineering Science, Yokohama National University, 79-5 Tokiwadai, Hotogaya-ku, Yokohama, 240-8501, Japan; Graduate School of Science, Kobe University, 1-1 Rokkodai-cho, Nada-ku, Kobe, 657-8501, Japan

**Keywords:** Amyloid fibril, prefibrillar intermediates, protein complex formation, amyloid inhibitor, SAXS, TEM, SEC, solid-state NMR

## Abstract

Amyloid fibrils are abnormal protein aggregates that relate to a large number of amyloidoses and neurodegenerative diseases. The oligomeric precursors, or prefibrillar intermediates, which emerge prior to the amyloid fibril formation, have been known to play a crucial role for the formation. Therefore, it is essential to elucidate the mechanisms of the structural development of the prefibrillar intermediates and ways to prevent its fibril formation. An insulin-derived peptide, insulin B chain, has been known for its stable accumulation of the prefibrillar intermediates. In this study, structural development of B chain prefibrillar intermediates was monitored by transmission electron microscopy and small-angle X-ray scattering combined with size exclusion chromatography and solid-state NMR spectroscopy to elucidate the stability and secondary structure. We further tracked its inhibition process by fibrinogen (Fg), which has been known to effectively prevent the amyloid fibril formation of B chain. We demonstrated that prefibrillar intermediates are wavy structures with low β-sheet content, growing in a multistep manner toward the nucleation for the amyloid fibril formation. In the presence of Fg, the formation of the prefibrillar intermediates slowed down by forming specific complexes. These observations suggest that the prefibrillar intermediates serve as reaction fields for the nucleation and its propagation for the amyloid fibril formation, whereas the inhibition of prefibrillar intermediate elongation by Fg is the significant factor to suppress the fibril formation. We propose that the obtained molecular picture could be a general inhibition mechanism of the amyloid fibril formation by the inhibitors.

## Introduction

Amyloid fibrils, which possess β-sheet-layered fibrous structures, are abnormal protein aggregates related to various kinds of diseases such as Alzheimer’s disease, Parkinson’s disease, and familial amyloidotic polyneuropathy.^1, 2^ As a general picture of the amyloid fibril formation, a nucleation-dependent polymerization has been proposed, in which formation of nuclei responsible for the fibril formation is a rate-limiting process. Once nuclei are formed, subsequent rapid elongation of fibrils from the nuclei occurs in a self-propagation manner.^3^ Unfolded or misfolded proteins are potential species to form nuclei via intermolecular interactions.^4^

Recently, it has been suggested that the nucleation occurs via on-pathway oligomeric species observed prior to the fibril formation, which we here refer to as prefibrillar intermediates.^3^ For example, one of the nucleation mechanisms describing this phenomenon is nuclear conformational conversion (NCC), which was firstly suggested in Sup35 study.^5^ In this mechanism, it is proposed that monomers first form oligomeric species, which accelerates the subsequent nucleation within the oligomers. Therefore, understanding the structural property of the prefibrillar intermediates is one of the essential ways to elucidate the mechanism of the amyloid fibril formation. Furthermore, there have been several reports showing that oligomers are more toxic than amyloid fibrils.^6, 7^ Thus, understanding the molecular property of prefibrillar intermediates would also be useful for developing therapeutic methods for preventing not only amyloid fibril formation but also their toxicities.

We have recently found an efficient and stable accumulation of prefibrillar intermediates of an insulin-derived peptide, insulin B chain (F^1^VNQHLCGSH^10^LVEALYLVCG^20^ERGFFYTPKT^30^; hereafter we simply refer to as B chain).^8^ It was revealed that the structural development occurs via two prefibrillar intermediates where the second one subsequently develops after the formation of the first one. Interestingly, it was revealed that the second prefibrillar intermediate plays a significant role for forming nuclei requisite for the amyloid fibril formation. These structural developments proceeded in a time scale from minutes to hours, and thus were able to be tracked by several spectroscopic and scattering techniques such as circular dichroism (CD) spectroscopy, nuclear magnetic resonance (NMR) spectroscopy, and dynamic light scattering (DLS).^8^ We observed that the formation of the first prefibrillar intermediate occurred < 30 min with the size development and partial secondary structure formation, followed by the formation of the second prefibrillar intermediate with further developments of the size and structure within ~500 min. This well-characterized B chain is thus a useful model system to study structural developments of the prefibrillar intermediates.

To prevent the structural development of prefibrillar intermediates which could lead to the amyloid fibril formation, various kinds of molecular chaperones play a crucial role for guiding correct protein folding, inhibiting amyloid fibril formation, and also inducing protein disaggregation in the intracellular environment of multicellular organisms.^9, 10^ Furthermore, there are an increasing number of reports of extracellular chaperones which inhibit amyloid fibril formation.^11^ These intracellular and extracellular chaperones are essential for maintaining protein homeostasis, i.e. proteostasis. In this context, we have found that a rod-shaped fibrinogen (Fg) with a molecular mass of ~340 kDa, which is known to be one of the potent extracellular chaperones,^11, 12^ inhibited the amyloid fibril formation of B chain by interacting with the prefibrillar intermediates. ^13^ It would thus be great interest to characterize the detailed process of the interaction which leads to the inhibition of the amyloid fibril formation. This could be helpful to understand how the chaperone molecule works for the prevention of the fibril formation.

In this paper, we report the structural development of the prefibrillar intermediates of B chain and the inhibition mechanism of the amyloid fibril formation by Fg using transmission electron microscopy (TEM) and small-angle X-ray scattering (SAXS). TEM is a widely-used tool to observe amyloid fibrils.^14^ SAXS is a powerful tool to investigate the structures of macromolecules in solutions,^15^ and has successfully tracked time-dependent structural development of prefibrillar intermediates or protofibrils of amyloid-prone proteins and peptides.^16–20^ First, using TEM, we show that B chain prefibrillar intermediates are a wavy rod-like structures accumulating prior to amyloid fibril formation, and the fibril formation is indeed prevented by Fg. Next, we track detailed structural development of the prefibrillar intermediates and its inhibition by Fg. Parameters such as the base diameter and length of the prefibrillar intermediates are obtained by combining SAXS with DLS, and compared with those obtained by TEM. Size exclusion chromatography (SEC) and solid-state NMR spectroscopy are also performed to obtain additional information on the structural stability of prefibrillar intermediates and the secondary structural changes on the formation of the amyloid fibrils. Based on these experimental results, the molecular mechanisms of the formation of the prefibrillar intermediates and its inhibition by Fg are discussed.

## Results

### Amyloid fibril formation via prefibrillar intermediates

To obtain the direct evidence of the formation of prefibrillar intermediates prior to the amyloid fibril formation, the fibril formation was traced by thioflavin T (ThT) fluorescence and TEM. The reaction was initiated by pH jump from ~11 (denatured monomer) to 8.7, and the sample was shaken at 1,200 rpm, which we term as the agitated condition. Figure 1A shows the time dependence of the ThT fluorescent intensity in the agitated condition at the B chain concentration of 1.4 mg/mL. The time dependence is approximated to a sigmoid curve, which is typical for the amyloid fibril formation. As shown in the inset of Figure 1A, slight intensity increase was observed in the up to 20 min. The TEM image taken at 20 min showed long, and thin wavy structures (I in Figure 1B). These are prefibrillar intermediates accumulated prior to the amyloid fibril formation. At around 30 min, the ThT intensity started to increase again, and at 40 min the co-existence of the prefibrillar intermediates and amyloid fibrils was confirmed (II in Figure 1B). At 50 min, the amyloid fibrils became dominant species (III in Figure 1B). At 80 min, the formation of the amyloid fibril was confirmed (IV in Figure 1B). These results strongly suggest that the amyloid fibril formation occurs via prefibrillar intermediates with long and thin filamentous morphology.

**Figure 1.**
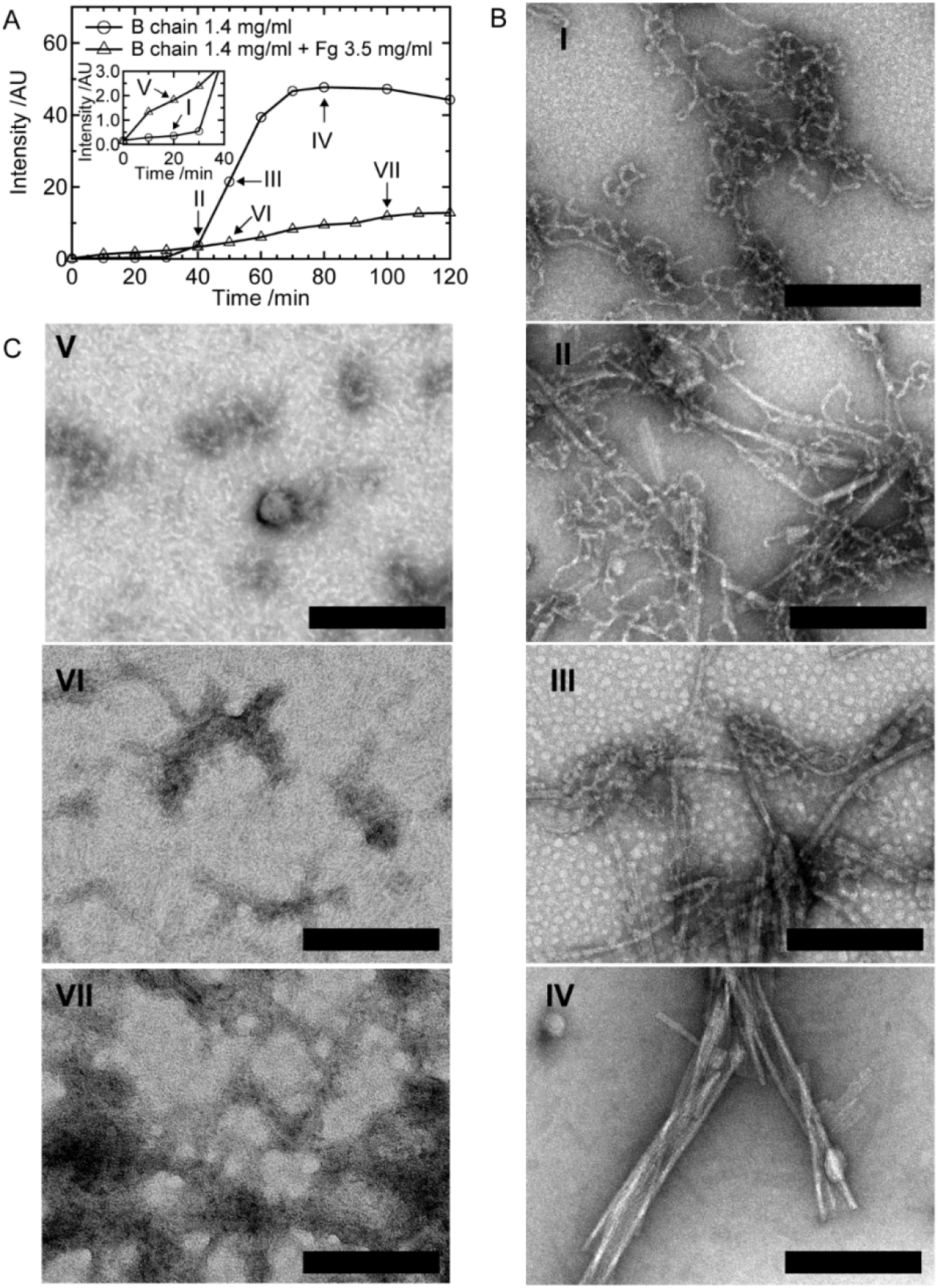
Amyloid fibril formation of B chain and its inhibition by Fg in the agitated condition. (A) Time course of ThT fluorescent intensity. The inset shows the magnified data in the early period. The Roman numerals correspond to times where TEM images shown in panel B or C were taken. (B and C) TEM images of the B chain amyloid fibril formation without (B) or with (C) Fg. The Roman numerals correspond to the times indicated in the panel A. The scale bars indicate 200 nm.

### Inhibition of amyloid fibril formation by Fg

To track how Fg inhibits the amyloid fibril formation, the time course of ThT fluorescent intensity was monitored in the presence of Fg. The experiment was performed at a Fg concentration of 3.5 mg/mL. At this concentration, the sigmoidal increase in ThT intensity observed in the absence of Fg was suppressed, suggesting that the interaction of Fg with B chain inhibits the amyloid fibril formation. The TEM image taken at 20 min (V in Figure 1C) shows that there were no thin and long structures, and instead, shorter fragments whose length are around 100 nm accumulated. At 50 min, or even at 100 min, apparent amyloid fibril formation was not confirmed (VI or VII in Figure 1C). The result demonstrates that Fg inhibits the amyloid fibril formation via inhibition of the elongation of prefibrillar intermediates.

### Prolonged existence of the prefibrillar intermediates in the quiescent condition

Figure 2A shows the time dependence of the ThT fluorescent intensity at 1.4 mg/mLB chain under the quiescent condition (circles), in which no agitation was applied. In this condition, the amyloid fibril formation became much slower than in the agitated condition (squares). ^8, 13^ The increase in the intensity was monotonic, and no sigmoidal increase as observed in the agitated condition was observed until 960 min (16 h). In agreement with this, wavy intermediates similar to those observed in the early stage in the agitated condition were identified as products (Figure 2B). This represents that the prefibrillar intermediates are stable as long as no agitation is applied. Hereafter, we focus on this quiescent condition to monitor the structural development of the prefibrillar intermediates and the interaction mechanism with Fg.

**Figure 2.**
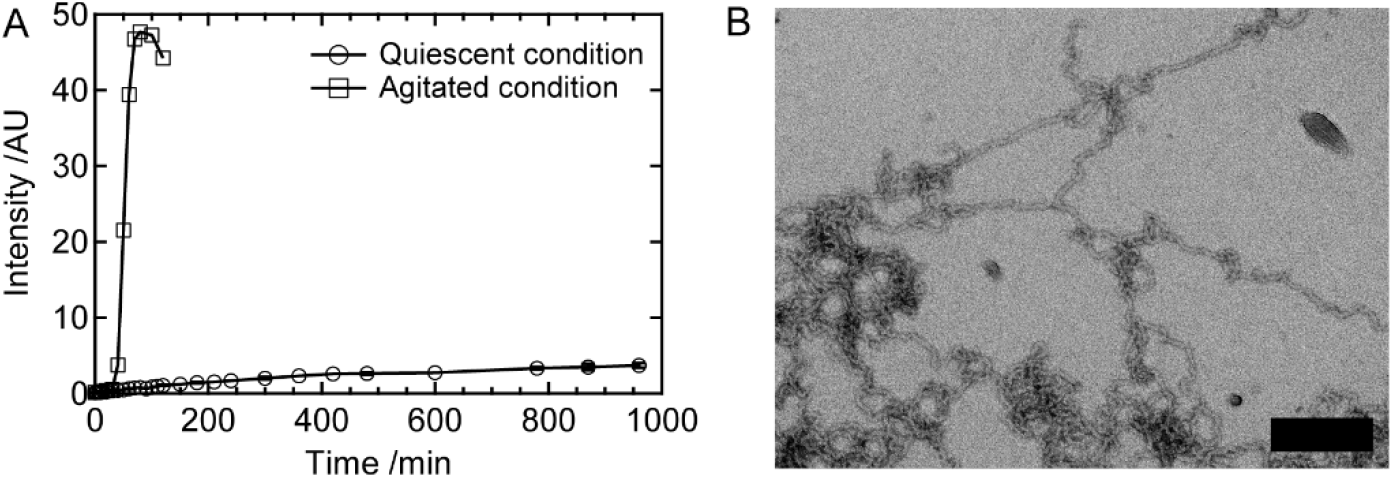
Accumulation of B chain prefibrillar intermediates in the quiescent condition. (A) Time course of ThT fluorescent intensity. For comparison, time course in the agitated condition (Figure A) is also shown. (B) A TEM image of the prefibrillar intermediates taken at 16 hours. The scale bar indicates 200 nm.

### Structural development of prefibrillar intermediates analyzed by SAXS and TEM

To obtain detailed structural insight into the prefibrillar intermediates, the structure development was traced by SAXS in the quiescent condition. Figure 3 shows SAXS time course of B chain at 1.4 mg/ml. As shown in Figure 3A, the scattering intensity gradually increased with time especially at low *q* region, which is quantitatively confirmed by the time-dependence of the intensity at *q* = 0.1 nm^-1^. The slope at the intermediate region of the SAXS profile was obtained by fitting with eq. 6 (lines in Figure 3A). As shown in Figure 3B, the value of the slope was almost constant at around −1 over the measurement time period. This indicates that the prefibrillar intermediates possess rod-like structures. We here note that ~60 % of B chain participates in the formation of the prefibrillar intermediates, and the rest remains monomer. [Yamamoto, 2018 #2] However, the scattering from monomer is negligible due to its small size compared to the intermediates.

**Figure 3.**
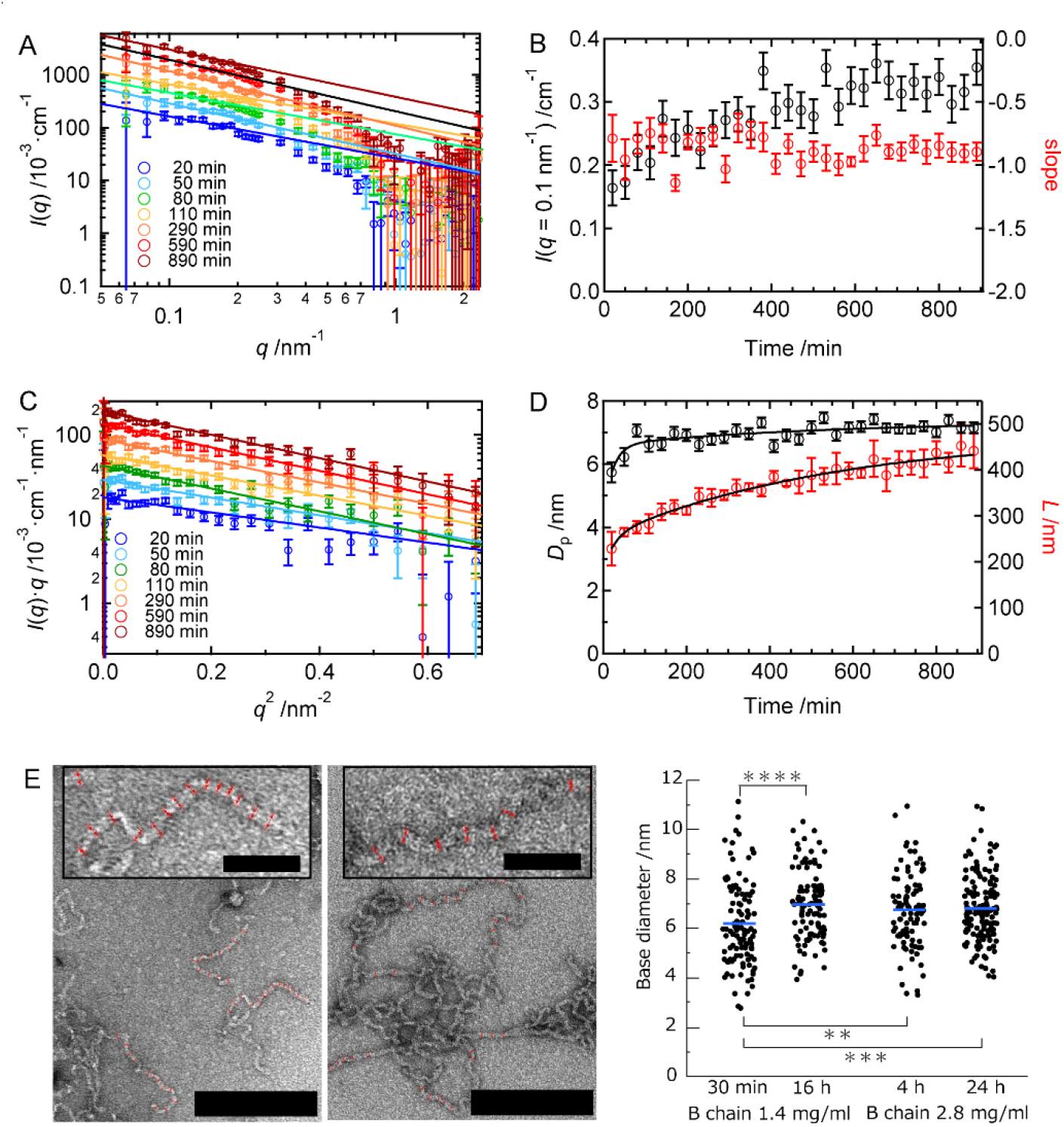
(A-D)Time-course of SAXS profiles of B chain alone at the concentration of 1.4 mg/ml. (A) The intensity vs *q* plot. The lines represent fitting results obtained by using eq. 6 at 0.07 ≤ *q* ≤ 0.15 nm^-1^. Each plot is arbitrary shifted for guiding eyes. (B) Time-dependence of the intensity at *q* = 0.1 nm^-1^ (black symbol) and the slope of the fitting lines shown in panel A (red symbol), respectively. Each plot is arbitrary shifted for guiding eyes. (C) The cross-section plot. The lines represent fitting results performed by using eq. 7 at 0.1 ≤ *q* ≤ 0.4 nm^-1^ to obtain the base diameter of gyration. (D) Time-dependence of the base diameter (black symbol) and the length. The solid lines represent the fitted curves. See detail in the main text. (E) Analysis of the base diameter at 1.4 mg/ml B chain using TEM images. TEM images used for analysis are shown. Left and right images are 30-min and 16-hours samples, respectively. The red lines with arrows are indicators used to measure the base diameter. The scale bar indicates 200 nm. The magnified parts are shown in the insets. The scale bar indicates 50 nm. The resultant statistics are shown in the right panel. Results of the higher concentration case, i.e. 2.8 mg/ml B chain are also shown. The blue lines indicate average values of each case. Differences considered to be significant are indicated as **; *P* < 0.01, ***; *P* < 0.005, and ****; *P* < 0.001, respectively. If *P* > 0.05, no significance was considered or no indicators are put in the panel.

The advantage of SAXS measurement is the ability to estimate the size of scattered particles. The base diameter of the rod of the prefibrillar intermediates, *D*_p_, was obtained by analyzing the cross-section plot using eqs. 7 and 8. Figures 3C and D show the experimental plot and the *D*_p_ values obtained from the regression. As shown in Figure 3D, *D*_p_ increased in a bi-exponential manner. This is consistent with the time dependence of DLS (Figure S1). Then, the time dependence of *D*_p_ obtained in this study was fitted using a bi-exponential equation below;

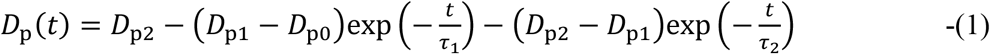

where *D*_p0_, *D*_p1_ and *D*_p2_ represent the base diamters of the initial, first, and second species where the development of a first species followed by a second one is characterized by *τ*_1_ and *τ*_2_, respectively. The resultant values were *D*_p0_ = 4.4 ± 1.6 nm, *D*_p1_ = 6.6 ± 0.2 nm, and *D*_p2_ = 7.4 ± 0.2 nm, respectively, with time constants of *τ*_1_ = 22 ± 16 min and *τ*_2_ = 560 ± 90 min (resultant value ± SD calculated by curve fitting). These time constants are similar to those obtained by CD spectra in the previous study, which were assigned as the development of the first prefibrillar intermediate (*τ*_1_ = 16 ± 2 min) followed by the second prefibrillar intermediate (*τ*_2_ = 530 ± 80 min).^8^

As seen in Figure 1B and 2B, the prefibrillar intermediates in TEM images possessed high potency to get entangled with each other during the sample preparation process, which prevented the quantitative evaluation of the length based on TEM images. We thus calculated the length based on eq. 10 using the value of *D*_p_ obtained by SAXS combined with the hydrodynamic diameter, *D*_h_, obtained from the DLS measurement (SI methods and Figure S1). As shown in Figure 3D, the increase in length also seems to be a bi-exponential manner. We then fitted the time dependence using the following bi-exponential function;

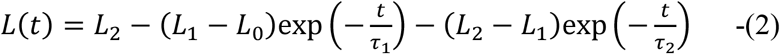

where *L*_0_, *L*_1_ and *L*_2_ represent the length of the initial, first, and second states where the developments from the initial to first state and from the first to second state are characterized by *τ*_1_ and *τ*_2_, respectively. The resultant values were *L*_0_ = 180 ± 70 nm, *L*_1_ = 250 ± 20 nm, and *L*_2_ = 480 ± 30 nm, with *τ*_1_ = 25 ± 30 min and *τ*_2_ = 550 ± 170 min (resultant value ± SD calculated by curve fitting). Similar to the case of the base diameter, these time constants are close to those obtained by CD spectroscopy in the previous study, although it was difficult to obtain the *τ*_1_ value accurately because the measurement interval was longer than that used for the CD analysis. These results demonstrate that the structural development of the prefibrillar intermediate is in a two-step manner; from first to second prefibrillar intermediates accompanying with increase in both the base diameter and length of the rod-like structures.

The structural development from the first to the second prefibrillar intermediate was further verified by measuring the base diameter of the first and second prefibrillar intermediates using TEM images in Figure S2 obtained at 30 min and 16 hours where the 1st and 2nd prefibrillar intermediates are the main components, respectively. Figure 3E shows the analyzed TEM images and resultant base diameters. The average base diameters of the first and second prefibrillar intermediate were 6.2 ± 2.9 nm and 7.0 ± 1.9 nm, respectively (average ± SD). T test was performed using these two quantities, and the resultant P-value was 0.00018, representing that the difference is significant. Furthermore, these average values coincided with those obtained by the SAXS analysis (6.6 ± 0.2 nm and 7.4 ± 0.2 nm, respectively). Therefore, the second prefibrillar intermediate is thicker than the first one in average. The difference in the absolute values in the TEM and SAXS results could be due to a partial coverage of edges of the prefibrillar intermediates by the dye in TEM images.

### Analyzing stability of the prefibrillar intermediates by SEC

We also performed SEC to compare the difference in stability of intermolecular interactions between the first and the second prefibrillar intermediates. We previously found that prefibrillar intermediates dissociate into monomers upon dilution,^13^ which we focused on for evaluating stability in this study. The samples incubated at 30 min and 16 hours, were used for the first and second prefibrillar intermediates, respectively. The results are shown in Figure 4A. The first prefibrillar intermediate possessed an elution peak around 11 ml (marked by *) and 17 ml (marked by †). TEM showed that short fragments as well as particles coexist at the ~11 ml elution fraction (Figure 4B). This fragmentation of the prefibrillar intermediates is consistent with the fact that they dissociate into monomers upon dilution. ^13^ Interestingly, some of the fragments accompanied particles in their termini (as indicated by arrows in Figure 4B). These particles could be monomers or oligomers that are dissociating form the prefibrillar intermediates. In the 17 ml elution fraction, particles around 10 nm were observed (Figure 4C, black arrows). This elution volume was close to that of 0.35 mg/ml B chain (black peak at 17.6 ml in Figure 4A), which possesses random-coiled monomer.^8^ Indeed, the diameter of this monomer (Figure 4D, black arrows) is similar to those observed in the 17 ml elution fraction. Therefore, the particles observed in the 17 ml elution fraction are supposed to be a random-colied B chain monomer.

**Figure 4.**
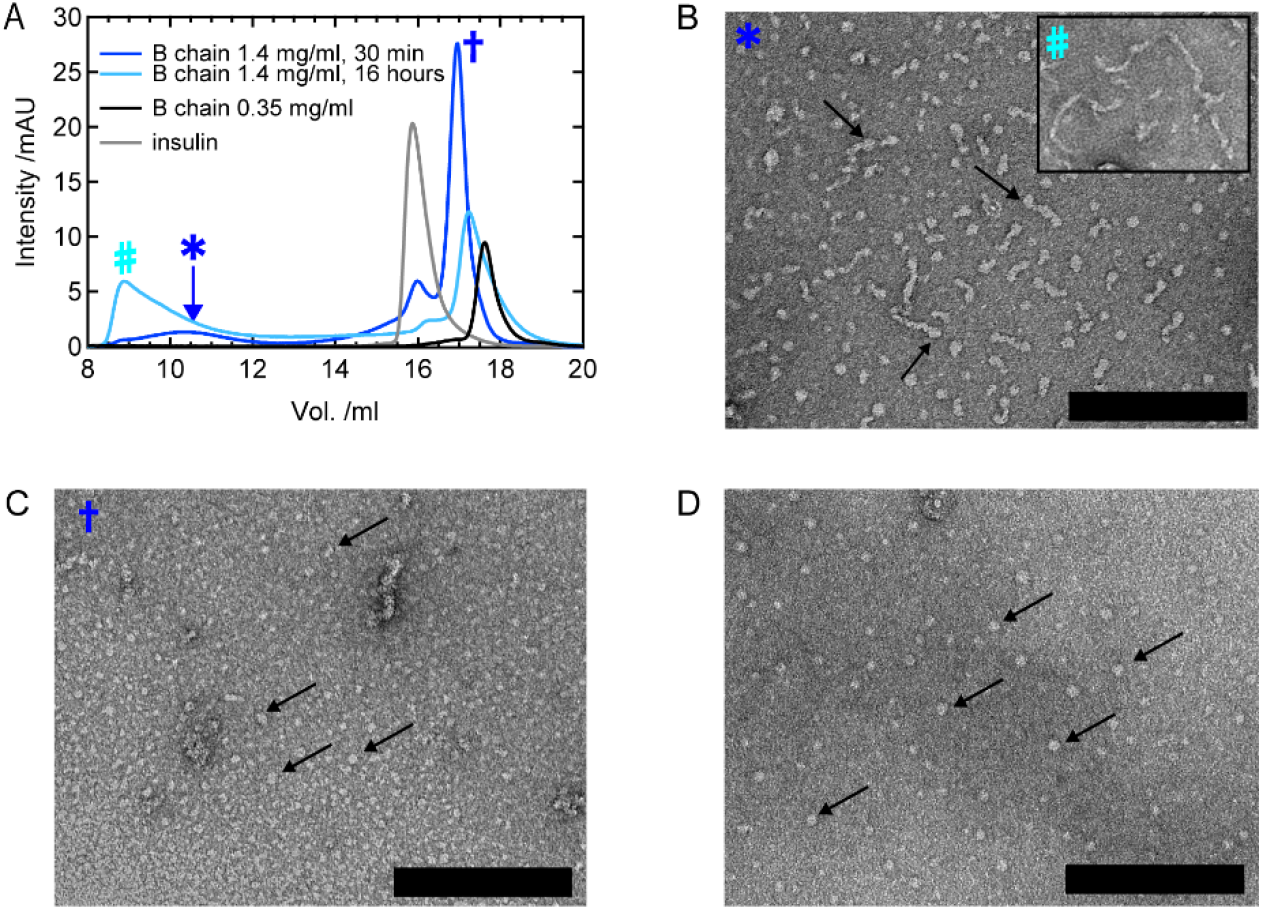
SEC and TEM analyses of the B chain prefibrillar intermediates at 1.4 mg/ml in the quiescent condition. (A) SEC charts of the prefibrillar intermediates accumulated at 30 min or 16 hours, respectively. As references, B chain monomer (0.35 mg/ml) and insulin (1 mg/ml) were also analyzed. The fractions for TEM were indicated by the symbols, which are used in panel B, C, and D. (B and C) TEM images of the 30-min sample at ~11 ml and 17 ml fractions, respectively. The inset in B shows a TEM image of the 16-hours sample. (D) A TEM image of 0.35 mg/ml B chain appearing at ~17.5 ml fraction. The scale bars in these TEM images indicate 200 nm.

In the case of the 16-hour sample, in which the second prefibrillar intermediate is the major component, it possessed a fraction at ~9 ml (marked by # in Figure 4A), which was larger and earlier than the peak of ~11 ml observed for the 30-min sample. In conjunction with this, the relative fraction volume at 17 ml of the 16-hours sample was smaller than that of the 30-min sample (Figure 4A). Some of the short fragments indeed showed longer lengths than that of the 30-min sample (Figure 4B, inset). These results indicate that the structure of the second prefibrillar intermediate is more stable than that of the first prefibrillar intermediate against dilution.

### Further structural development of the second prefibrillar intermediates at a higher B chain concentration

A further structural development that might subsequently occur after the formation of the second prefibrillar intermediate will contain structural information coupled to the amyloid fibril formation. To observe this, time-dependence at the twice concentration of 1.4 mg/ml, i.e. 2.8 mg/ml, was monitored by SAXS. Like the case of 1.4 mg/ml, the intensity increased as a function of time with maintaining a rod-like structure (Figure S3A and B). When the base diameter was estimated by fitting the cross-section plot (Figure S3C) using eqs. 7 and 8, it initially increased until ~200 min in a single exponential manner, and then reached plateau (black line in Figure 5A). The time course was then analyzed using a single exponential function;

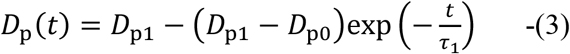

where *D*_p0_ and *D*_p1_ represent initial and final diameter of the rod, respectively, whose time change is characterized by the time constant *τ*_1_. The obtained values were *D*_p0_ = 6.4 ± 0.2 nm and *D*_p1_ = 7.4 ± 0.1 nm with *τ*_1_ = 112 ± 25 min (resultant value ± SD calculated by curve fitting). The values of *D*_p0_ and *D*_p1_ are close to those of the first and the second prefibrillar intermediates at 1.4 mg/ml B chain, respectively, indicating that this process corresponds to the formation of the second prefibrillar intermediate from the first one. The spontaneous formation of the first prefibrillar intermediate is supported by solution ^1^H-NMR measurements where signal intensities quickly decreased within 10 min (Figure S4). The analysis of TEM images also showed that the average value of the base diameter at 4 hours (6.8 ± 2.3 nm, average ± SD) at 24 hours (6.8 ± 1.8 nm, average ± SD) were similar to that of the second prefibrillar intermediate at B chain of 1.4 mg/ml (Figure 3E).

**Figure 5.**
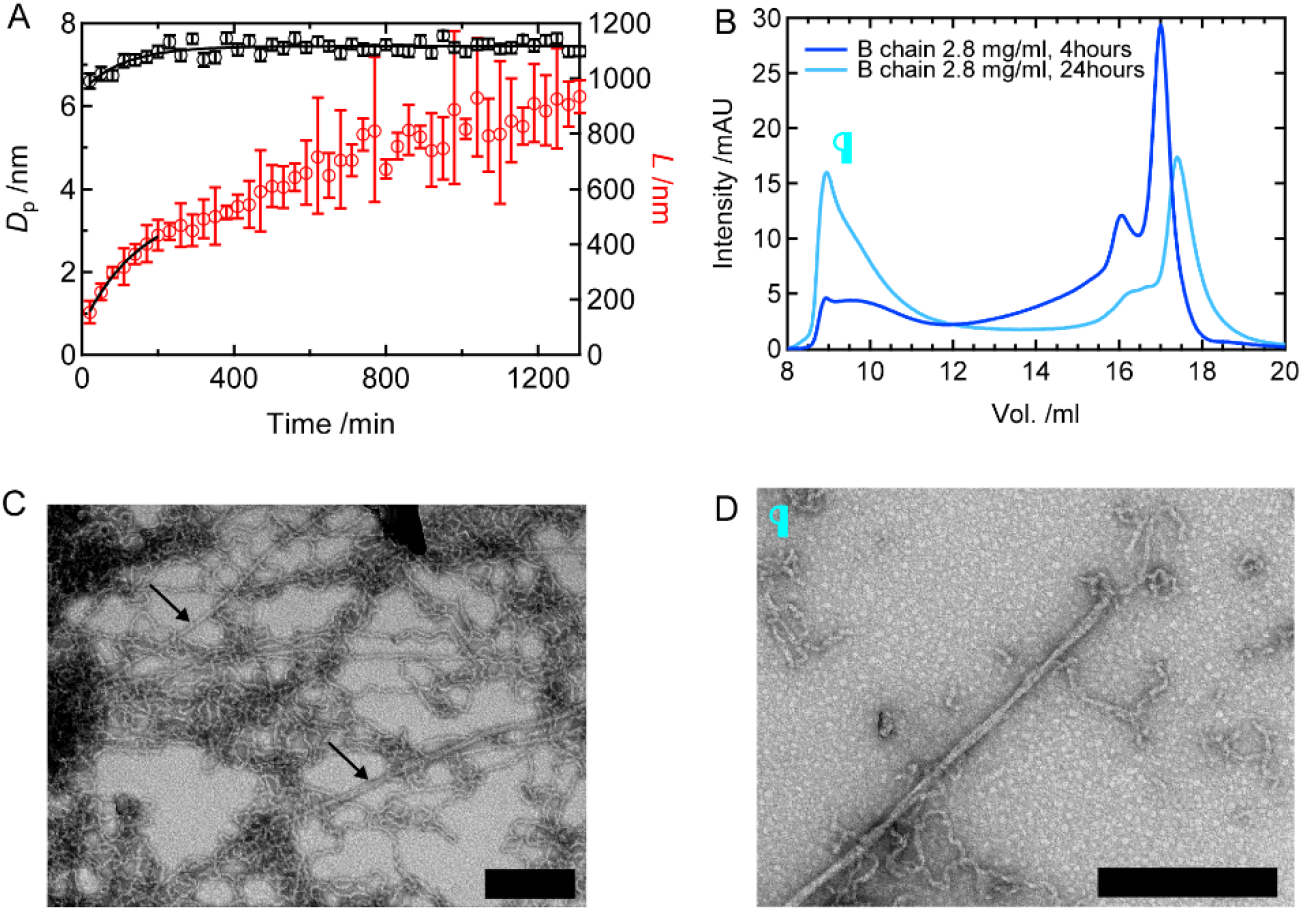
Characterization of prefibrillar intermediates of B chain at 2.8 mg/ml by SAXS, TEM, and SEC. (A) Time dependences of the base diameter and length obtained from SAXS combined with DLS. (B) SEC results. Symbols indicate elution fractions where TEM images were taken. (C) A TEM image taken at 24 hours. The black arrow indicates an amyloid fibril. The scale bar indicates 200 nm. (D) A TEM image of ~9 ml fraction at 24 hours. The scale bar indicates 200 nm.

The length was also calculated in the same manner as the lower concentration case using eq. 10 (red plots in Figure 5A). The length first steeply increased up to ~200 min, followed by an almost linear increase. We thus performed curve fitting for the initial steep increase using a single exponential function as follows;

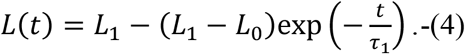

We obtained *L*_0_ = 103 ± 17 nm, *L*_1_ = 560 ± 70 nm, respectively, with *τ*_1_ = 157 ± 44 min (resultant value ± SD calculated by curve fitting). The fitting range was set until 200 min as this range could fairly be described by the single exponential function. The value of *L*_1_ is similar to *L*_2_ obtained at 1.4 mg/ml which was assigned as the length of the second prefibrillar intermediate (480 ± 30 nm), which reinforces the idea that the formation of the first prefibrillar intermediate spontaneously occurred within 20 min. After ~200 min, an additional linear increase in length continued to occur at least up to 1 day while the base diameter was kept constant, suggesting an additional linear elongation process of the second prefibrillar intermediate. Furthermore, the relative volume fraction in the SEC elution at ~9 ml increased compared to that of 4 hours (Figure 6C), which indicates that the structure of the second prefibrillar intermediate became more stable after the additional elongation. Interestingly, the TEM showed that a few amyloid fibrils emerged within the background of a large number of prefibrillar intermediates (Figure 5C, black arrows). The SEC analysis also revealed that long amyloid fibrils existed together with the prefibrillar intermediates at the ~9 ml elution (Figure 5D). These results imply that the amyloid fibril formation is triggered upon the stabilization of the second prefibrillar intermediate due to further elongation.

**Figure 6.**
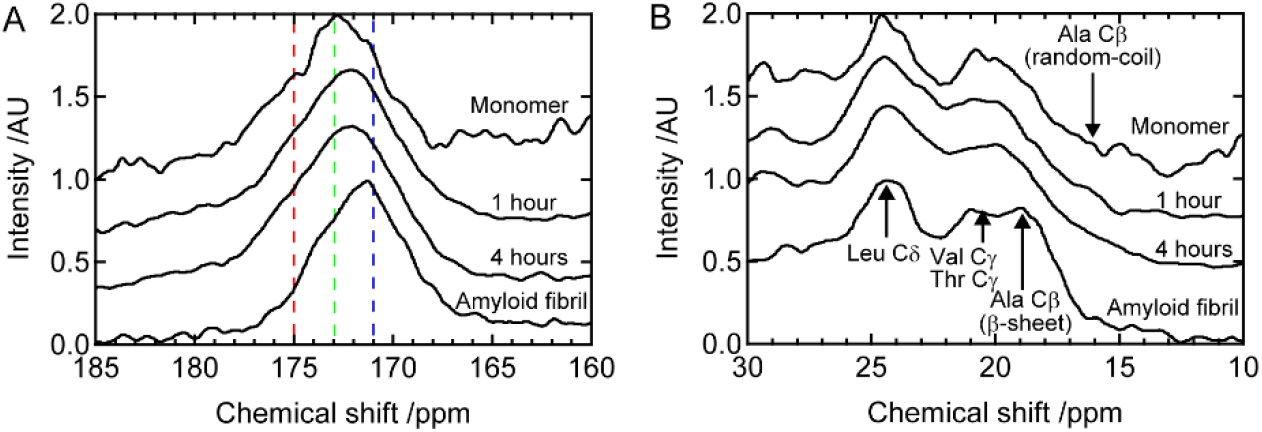
Solid-state ^13^C CP-MAS spectra of B chain alone in the lyophilized powder. (A) The main-chain carbonyl carbon region. Typical chemical shifts of an α-helix, random-coil, or β-sheet structure is indicated by the red, green, or blue dashed line. (B) The methyl/methylene carbon region. “1 hour” and “4 hours” indicates samples prepared 1 hour or 4 hours after the initiation of the fibril formation reaction. “monomer” means the monomeric state, which was prepared at the concentration of 0.35 mg/ml and pH 8.7. “amyloid fibril” indicate the amyloid fibril sample. To prepare this sample, B chain of 1.4 mg/ml at pH 8.7 was incubated with spinning at 1,200 rpm at 25 °C.

### Secondary structural changes of prefibrillar intermediates observed by ^13^C solid-state NMR

To obtain more detailed information concerning the structural developments of the prefibrillar intermediates, solid-state ^13^C NMR measurements were performed. The technique can detect signals from large molecules such as prefibrillar intermediates and amyloid fibrils which are not visible by solution-state NMR. We prepared two samples for monitoring formation of the prefibrillar intermediates; one is a powder sample lyophilized 1 hour after the initiation of the formation of the prefibrillar intermediates. This sample mainly contains the first prefibrillar intermediate where the content of the first and the second prefibrillar intermediates are ~90 % and ~ 10 %, respectively.^8^ The other is a powder sample lyophilized 4 hours after the initiation of the reaction where the amount of the second prefibrillar intermediate is expected to increase. We also prepared a lyophilized powder sample of B chain at the concentration of 0.35 mg/ml, which possesses a monomeric random-coil structure,^8^ and amyloid fibrils as references.

We obtained solid-state NMR spectra at the main-chain carbonyl region and side-chain methyl/methylene carbon region. In cross polarization-magic angle spinning (CP-MAS) NMR spectra of the lyophilized power state, all the carbons are detectable due to the low molecular mobility. Figure 6A shows ^13^C CP-MAS NMR spectra of the main-chain carbonyl carbons (160-185 ppm), which contain information of secondary structures. An α-helix structure typically shows their chemical shifts around 175 ppm (red dashed line), whereas that of the β-sheet structure is around 171 ppm (blue dashed line).^21^ The center of the spectrum in the monomeric state was around 172.5 ppm, which is near an averaged values of typical random-coil states (green dashed line).^22^ In contrast, the amyloid-fibril sample possessed a peak around 171 ppm, representing formation of the β-sheet structure.

The peak positions of the prefibrillar intermediates prepared 1 hour and 4 hours after the reaction started were at around 172 ppm, which is between those of the monomeric and amyloid-fibril states. The result strongly suggests that the prefibrillar intermediates form β-sheet structure partially. The β-sheet contents of the samples prepared 1 hour or 4 hours after the initiation of the reaction were 26.5 ± 3.2 % or 25.1 ± 3.3 %, respectively, as calculated by a spectral deconvolution analysis (Figure S5A). The fractions of the β-sheet content are much smaller than the populations of the prefibrillar intermediates obtained by ^1^H solution NMR (~60 %)^8^, and thus suggesting that only partial β-sheet formation occurs within a B chain molecule in the prefibrillar intermediates. No apparent difference in the β-sheet content within the error range was obtained between the prefibrillar intermediates at 1 hours and 4 hours, which is probably due to slight increase in the β-sheet content between these two times.

Figure 6B shows ^13^C CP-MAS spectra in the methyl/methylene carbon region (10-30 ppm). In B chain, eight Leu δ-carbons, six Val γ-carbons, two Thr γ-carbons, and one Ala β-carbon are included as methyl carbons. In all the sample conditions, several components were observed in the CP-MAS spectra. The peak at around 24 ppm is assigned as Leu δ-carbons.^21^ Signals at around 20-21 ppm are assigned as Val γ-carbons and Thr γ-carbons.^21^ Gln β-carbon and Glu β-carbon would also appear at around 25 ppm if these residues formed α-helix structures.^21^ However, the spectral analysis in the main-chain carbonyl region suggested that the samples do not contain α-helix structures (Figure 6A). In addition, only one Gln Cγ and one Glu Cγ are existing and thus the contribution of these residues is excluded from the NMR analysis at around 25 ppm. All the samples possess similar spectral features only with minor differences in the spectral region at 20-25 ppm, implying small secondary structure differences around the methyl groups among the different samples.

It has been reported that a signal of an Ala methyl carbon at around 20 ppm indicates the β-sheet structure, whereas that at around 16 ppm implies the random-coil state.^21^ B chain only possesses one Ala residue at 14th position, and thus structural information around this residue can be obtained by focusing on this signal. In the CP-MAS spectra, a broad component was observed at around 16 ppm in the monomer state, consistent with the fact that the structure is in a random-coil state with highly structural inhomogeneity. On the contrary, in the amyloid fibril state, the broad component around 16 ppm disappeared and a sharp peak around 19 ppm appeared instead. Interestingly, the spectra of the prefibrillar intermediates showed a broad component at 16 ppm as in the monomer state. This indicates that, while the middle region around the Ala14 residue of the B chain polypeptide is in a random-coil state possessing no specific structures in the prefibrillar state, it changes to the β-sheet structure upon the amyloid fibril formation. DD-MAS NMR spectra also share fundamentally similar spectral feature to those of CP-MAS spectra (Figure S5B), probably because methyl carbons show flipping motion about C3 axis, and thus strongly appear in DD-MAS.^23^

### Inhibition process of the elongation of the prefibrillar intermediates by Fg monitored using SAXS, TEM, and SEC

Figure 7 shows the time course of the SAXS profile of B chain at 1.4 mg/ml incubated with Fg of 3.5 mg/ml, under which fibril formation of B chain is fully suppressed. ^13^ We could track gradual increase in SAXS scattering intensity (Figure S6A) with the slope in the intermediate region at around −1 constantly (Figure S6A and B), representing a rod-like structure. The base diameter obtained by fitting the cross-section plot using eqs. 7 and 8 (Figure S6C) was time-independent within the error range (Figure 7A, top). The mean base diameter was 14.2 ± 0.2 nm (resultant value ± SD obtained by averaging all data points), which is larger compared to that of the first or second prefibrillar intermediate of Fg-free conditions (Figure 3D, ~6.6 nm or ~7.4 nm, respectively), indicating that Fg binds to the surface of the prefibrillar intermediates. The length of Fg-B chain complex was calculated using eq. 10. As shown in the top panel of Figure 7A, the resultant value of *L* monotonically increased in a single exponential manner approaching ~100 nm, demonstrating that the elongation of the prefibrillar intermediates is suppressed compared to the Fg-free condition (Figure 3D). Its time dependency was thus analyzed using eq. 4, and the resultant values were *L*_0_ = 69 ± 3 nm and *L*_1_ = 115 ± 2 nm with *τ*_1_ = 200 ± 30 min. A TEM image taken at the point where the reaction completed also indicated the presence of shorter fragments (Figure 7B) compared to a Fg-free condition (Figure 2B). Boundaries of particles are more or less ambiguous compared to those of the Fg-free conditions, which might suggest that Fg molecules surrounding the prefibrillar intermediates have tendency to be over-stained by uranyl acetate and aggregates during the sample preparation process as seen in Fg-only sample (Figure 7B, inset). The value of *τ*_1_ is much slower than the time constant of the formation of the first prefibrillar intermediate in the presence of Fg obtained by CD spectroscopy (21 ± 3 min),^13^ and also a rapid increase in *D*_h_ in the early time observed by DLS (Figure S1, *τ*_1_ =15 ± 3 min). These results suggest that the formation of the Fg-B chain complex follows the formation of the first prefibrillar intermediate.

**Figure 7.**
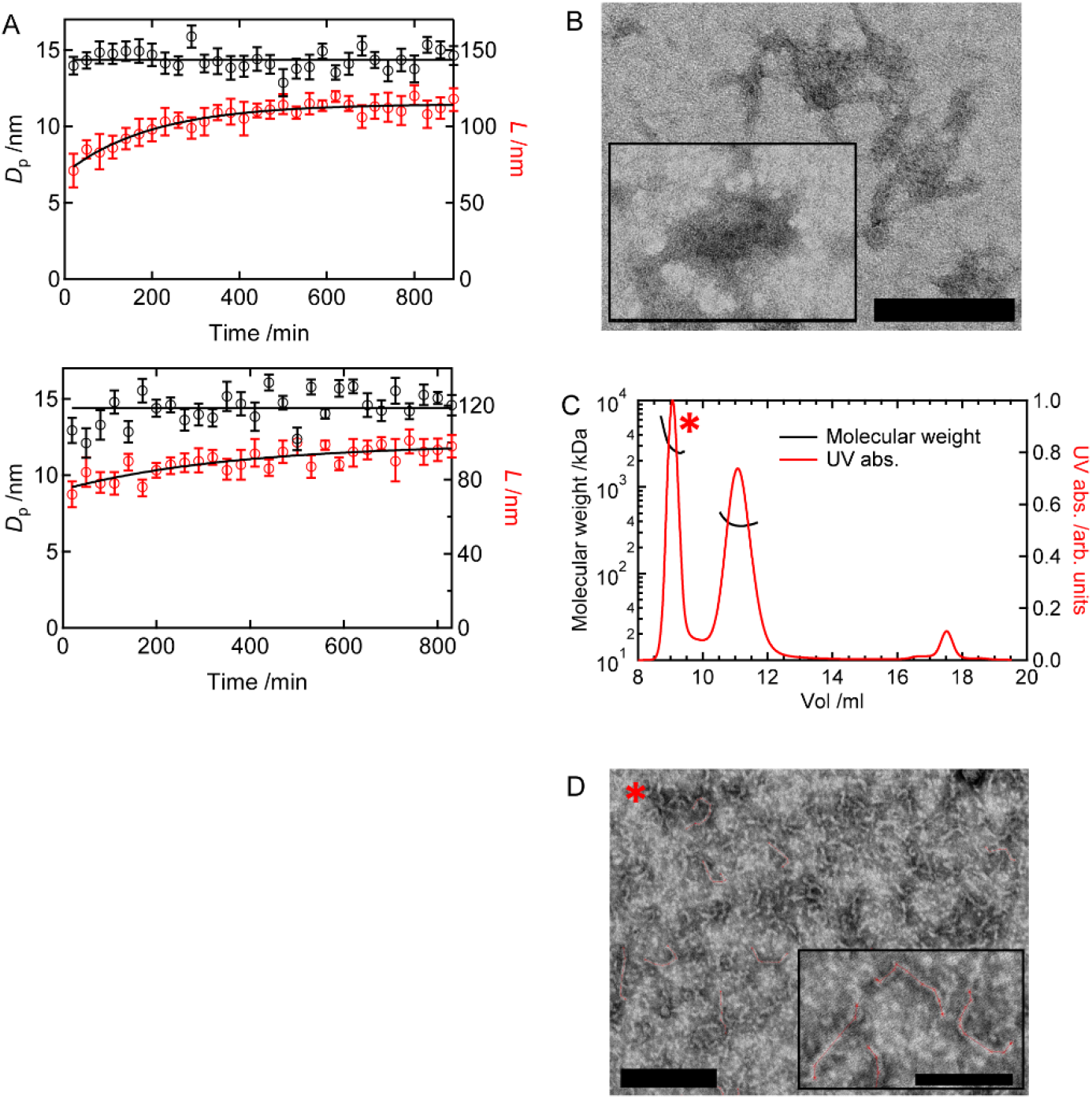
Characterization of the interaction of Fg with B chain prefibrillar intermediates by SAXS, SEC, and TEM. (A) Time dependence of the base diameter and length calculated by SAXS combined with DLS of 1.4 mg/ml B chain with 3.5 mg/ml Fg (top) and 2.8 mg/ml B chain with 7.0 mg/ml Fg (bottom), respectively. (B) A TEM image of B chain 1.4 mg/ml with Fg 3.5 mg/ml incubated for 19 h. The inset shows a TEM image of Fg only (3.5 mg/ml). (C) A SEC result at B chain of 1.4 mg/ml incubated with Fg of 3.5 mg/ml for 18 hours. The symbol indicates a fraction where the TEM image shown in panel D was taken. (D) The TEM image of ~9 ml SEC fraction of 1.4 mg/ml B chain with 3.5 mg/ml Fg incubated for 18 h. The scale bar indicates 200 nm. An example of the length analysis is enlarged in the inset where the scale bar indicates 100 nm. The red lines show traces used for the length estimation.

The complex formation between the prefibrillar intermediates and Fg was also confirmed using SEC. As shown in Figure 7C, one elution fraction appeared at ~9 ml (marked by *) besides another one at ~11 ml. The former one is considered to be the complex whereas the latter is unbound Fg (Figure S6D). To confirm this, their molecular weights were investigated using multi-angle light scattering (MALS) combined with SEC. As shown in Figure 7C, the molecular weight at ~11 ml elution fraction is ~350 kDa, which is consistent with that of Fg (340 kDa). On the other hand, the molecular weight of ~9 ml elution fraction ranged ~2,500 to ~6,500 kDa, which is much larger than that of Fg, indicating the elution of the complex. The wide distribution of the molecular weight might be due to partial dissociation of Fg molecules from the complex upon elution, although details remain unknown. TEM also confirmed fragmented prefibrillar intermediate-like structures with ambiguous boundaries in this elution fraction (Figure 7D). The average length was calculated by measuring those of isolated fragments (Figure 7D, inset), resulting in 101 ± 16 nm. The value is consistent with that obtained by SAXS (Figure 7A, 115 ± 2 nm).

Fg could also have potential to suppress further elongation from the second prefibrillar intermediates observed at the B chain concentration of 2.8 mg/ml (Figure 5A). To check this, SAXS profile was monitored in the time-dependent manner at the twice-higher concentration condition of the complex, i.e. 2.8 mg/ml B chain and 7.0 mg/ml Fg. The SAXS intensity gradually increased as a function of time with the slope at around −1 (Figure S7A, B), representing that the complex possesses a rod-like structure. The base diameter of the rod structure was obtained by fitting the cross-section plot using eqs. 7 and 8 (Figure S7C). Similar to the lower concentration case, the base diameter was time-independent within the error range (Figure 7A, bottom). The mean base diameter was thus obtained by fitting the data using an intercept. The mean base diameter was 14.4 ± 0.2 nm (resultant value ± SD obtained by averaging all data points). Interestingly, the value is almost identical to that obtained in the lower concentration condition (Figure 7A top; 14.2 ± 0.2 nm at B chain 1.4 mg/ml with Fg 3.5 mg/ml). The length was also calculated using eq. 10, which increased in a single exponential manner (Figure 7A, bottom). The time-dependence was thus fitted using eq. 10, resulting in *L*_0_ = 75 ± 3 nm and *L*_1_ = 99 ± 5 nm with *τ*_1_ = 370 ± 200 min, respectively (resultant value ± SD calculated by curve fitting). As with the case of the base diameter, the value of *L*_1_ is also close to that of the lower concentration condition (Figure 7A top; 115 ± 2 nm). Furthermore, SEC showed a similar elution fraction at ~9 ml to that of the lower concentration condition (Figure S7D, marked by §), and similar fragmentated prefibrillar intermediates were also observed by TEM (Figure S7E).

## Discussion

In this study, we scrutinized the molecular mechanism of the structural formation of the prefibrillar intermediates and the interaction between the prefibrillar intermediates and Fg, by using TEM, SAXS, SEC, and solid-state NMR. Based on the results, we discuss the mechanism of the amyloid fibril formation via prefibrillar intermediates as well as the inhibition mechanism of the fibril formation by Fg.

Figure 8 shows a schematic picture representing the amyloid fibril formation and its inhibition by Fg suggested in this study. It was revealed that the first and second prefibrillar intermediates have a wavy structure that can be approximated as a rod-like shape. The first prefibrillar intermediate, which is observed at the B chain concentration of 1.4 mg/ml, possesses the base diameter and length of ~6.6 nm and ~250 nm, respectively (Figure 3D). The second prefibrillar intermediate is a thicker and longer noodle-like structure whose base diameter and length are ~ 7.4 nm and 500 nm, respectively (Figures 3D and 5A). After the formation of the second prefibrillar intermediate, further elongation is observed at the B chain concentration of 2.8 mg/ml (Figure 5A). In this process, nucleation and its propagation tend to occur, leading to the formation of amyloid fibrils (Figure 5C). In the presence of Fg, a complex with the base diameter and length of ~14 nm and ~70 nm, respectively, are rapidly formed, which slowly develops to the longer size (~110 nm) while maintaining the base diameter (Figure 7A). The formation of the specific complex is considered to play a role to inhibit the amyloid fibril formation.

**Figure 8.**
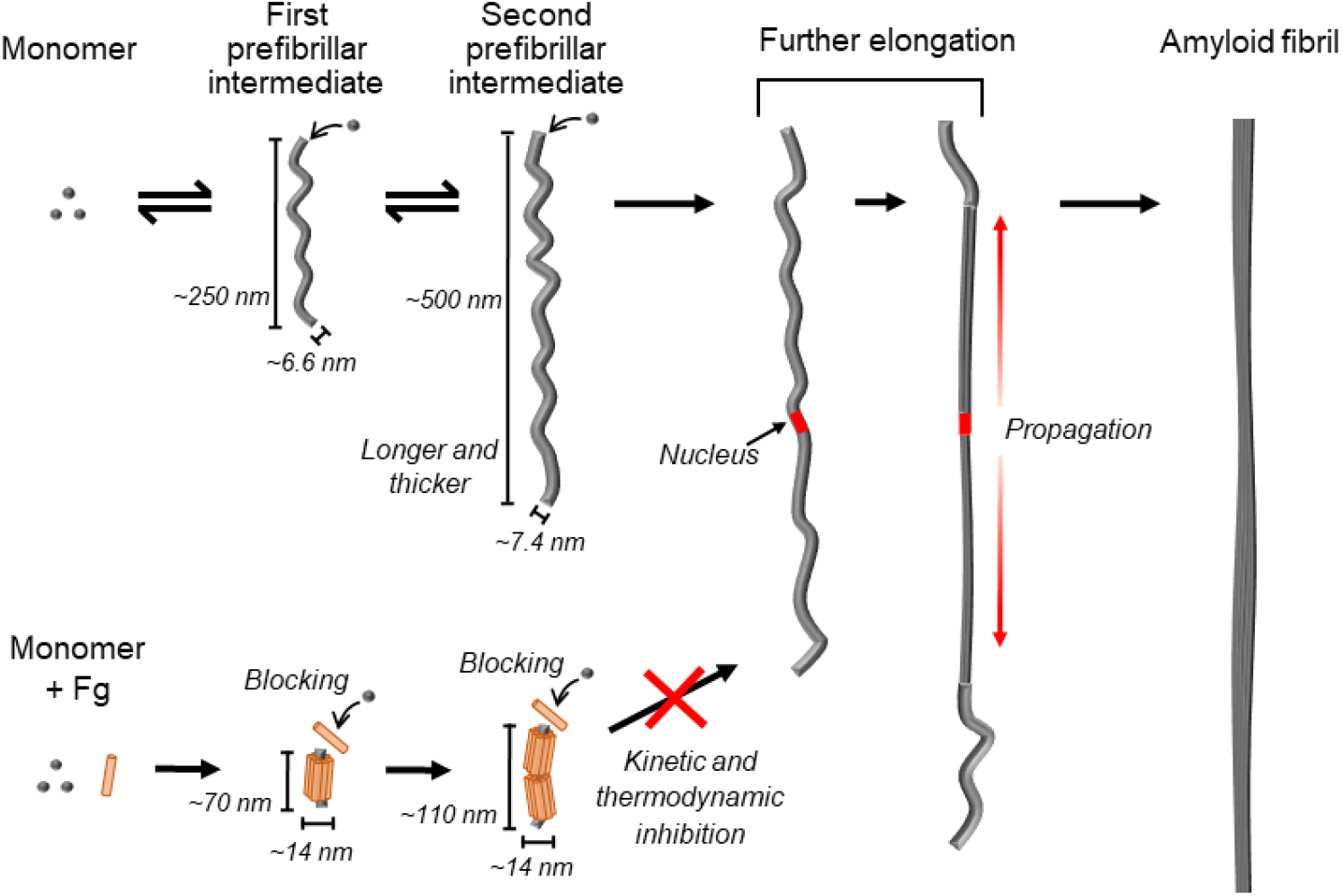
A proposed schematic picture of the structural developments of B chain prefibrillar intermediates and their interaction with Fg. The upper chart is the structural development of monomeric B chain to the first prefibrillar intermediate the second prefibrillar intermediate, which further elongate where nucleation and its propagation could occur, resulting in the formation of the amyloid fibril. The lower chart is the situation in the presence of Fg. Fg interacts with the surface of the prefibrillar intermediate, which results in the formation of the stable complex. Fg might be blocking interaction of monomers or oligomers required for the elongation of the prefibrillar intermediates. As a result of these interactions, the amyloid fibril formation could be prevented. See detail in the main text.

### The role of the prefibrillar intermediates for the amyloid fibril formation

The prefibrillar intermediates observed by TEM had wavy morphology (Figures 2B, 3E, and 5C) and their elution by SEC showed low stability leading to the reversible dissociation of monomers from the prefibrillar intermediates (Figure 4B). The solid-state NMR measurement revealed that the amount of the β-sheet structure at 1.4 mg/ml B chain was 26.5 ± 3.2 % and 25.1 ± 3.3 % at 1 and 4 hours, respectively (Figure 6A and S5A). With a consideration that the fraction of the residual random-coil monomers was ~40 %, ^8^ it is implied that 44.2 ± 5.3 % and 41.8 ± 5.5 % of the whole peptide sequence possesses β-sheet structure at 1 and 4 hours, respectively. The low β-sheet content is reminiscent of weak intermolecular interactions within the prefibrillar intermediates. This would bring flexibility of the structures, as observed in the large distribution of the base diameters of the prefibrillar intermediates (Figure 3E). The wavy morphology of the prefibrillar intermediates might originate from the flexibility of the structure. These flexible and unstable properties of the prefibrillar intermediates are in clear contrast to the amyloid fibril where rigid and stable β-sheet structures are formed (Figure 5D and 6A).

Regarding the difference in structure between the first and the second prefibrillar intermediate, the second one has higher stability than the first one as shown by the SEC result (Figure 4A). The certain difference in the base diameter was also observed from the SAXS profiles and TEM images (Figure 3D and E). The difference in the secondary structure content, on the other hand, was not obvious between the first and second prefibrillar intermediate from the solid-state NMR spectra (Figure 6 and S5). These observations indicate that structural rearrangements are occurring inside the prefibrillar intermediate without apparent changes in the secondary structure. As the CP-MAS NMR spectra suggested, β-sheet structure around the 14th Ala residue was formed in the fibril form, and not in the prefibrillar intermediates (Figure 6B). Ala14 is located in the middle of the peptide sequence showing high propensity to form steric zipper (i.e., VEALYL).^24^ A maturation of a similar region was also observed in a fibrillation process from an intermediate of insulin in a high salt concentration condition.^25^ The present result together with that of the insulin fibril implies that the β-strand formation around the Ala residue possessing high fibrillation propensity is critical for the intermolecular interaction required for the maturation of the amyloid fibril formation. The structural conversion around the Ala residue could occur during the further elongation process from the second prefibrillar intermediate, which might play a role as a nucleation required for the fibril formation.

In the further elongation process of the second prefibrillar intermediate observed at 2.8 mg/ml B chain, further stabilization was observed (Figure 5B). Given that the amyloid fibril formation becomes to occur more easily as the elongation of the prefibrillar intermediate proceeds (Figure 5C), structural rearrangements accompanied by the elongation would be important for the amyloid fibril formation. There are several reports showing that the elongation of such rod-like structures as the prefibrillar intermediates of B chain plays a key role for the amyloid fibril formation. For example, Vestergaard et al. reported a SAXS study of an amyloid fibril formation of insulin.^19^ They proposed that an insulin helical hexamer formed in a beads-on-string manner constructs protofilaments that could finally become the amyloid fibril. They concluded that this hexamer is the structural nucleus for the amyloid fibril formation. Oliveira et al. reported a SAXS study on the amyloid fibril formation of glucagon.^18^ They proposed that prior to the amyloid fibril formation, glucagon forms rod-like intermediate states. The radius and length of the intermediate states develop as a function of time, which finally leads to the formation of matured amyloid fibrils. Based on these observations or reports, they concluded that the intermediates are on-pathway species that easily convert to mature amyloid fibrils. Taken together, formation of rod-like prefibrillar intermediates would function as a reaction field where the nucleation and following propagation efficiently occurs (“Further elongation” step in Figure 8). This propagation is supported by the TEM image in Figure 6D, where the terminal of the amyloid fibril appears to remain filamentous.

### The inhibition mechanism of the amyloid fibril formation by Fg

We revealed that the base diameter of the first and second prefibrillar intermediates in the Fg-free condition are ~6.6 or ~7.4 nm, respectively (Figures 3D and 5A), which increased to ~14 nm in the presence of Fg (Figure 7A). These results are consistent with a previous study where it was proposed that seven Fg molecules whose base diameter is ~5 nm align around the circumference of the prefibrillar intermediates along with its long axis.^13^ The proposed model of the complex structure has been reconfirmed in this study, and the detailed information on the structural development of the complex was obtained by the time-dependent tracking of SAXS and TEM.

Given the fact that the initial and final lengths of the complex are 70-75 nm and 100-110 nm (Figure 7A), respectively, and that of Fg is ~50 nm, one or two repeats of the alignment of seven Fg molecules are expected to incorporate in the interaction with the intermediates (Figure 8, bottom). This fact that the length of the complex is the integral multiples of Fg, it is speculated that the Fg molecule plays a role to regulate the length of the prefibrillar intermediates. In our previous study, the molar ratio of B chain per Fg required for the complete inhibition is estimated to be ~30.^13^ Using this value, the molecular mass of the complex with the length of ~100 nm could be ~6,200 kDa (see detail in SI methods). This value is consistent with the molecular mass obtained by MALS (Figure 7C).

The length of the complex (i.e., ~70 nm or ~110 nm) is much shorter compared to those of the first and second prefibrillar intermediates in the Fg-free condition (~250 nm or ~500 nm, respectively, Figures 3D and 5A), representing that Fg inhibits the elongation of the prefibrillar intermediates. A hint to understand the mechanism of this inhibition is found in the concentration-dependent experiment; the time constant of the elongation of the prefibrillar intermediates at B chain 2.8 mg/ml and Fg 7.0 mg/ml was larger than that at the half-concentration (Figure 7A, top and bottom) in spite of the higher concentrations of the constituent molecules, and thus some concentration-dependent kinetic driving force works for the inhibition. Possibly, free Fg molecules are dynamically interacting with the terminals of the prefibrillar intermediates, which would prevent association of B-chain monomers with the terminals required for the further elongation of the prefibrillar intermediates (“*Blocking*” in Figure 8).

The time constant of the formation of the second prefibrillar intermediate obtained by far-UV CD spectroscopy in the presence of Fg (1,200 ± 200 min at B chain and Fg of 1.4 mg/ml and 3.5 mg/ml, respectively)^13^ is much longer than that in Fg-free condition (530 ± 80 min)^8^. This fact implies that the inhibition of elongation by Fg contributes to suppresses the structural conversion from the first to second prefibrillar intermediate. Nevertheless, interestingly, it has also been known that the second prefibrillar intermediate was stabilized in the presence of Fg with respect to thermodynamics.^13^ This thermodynamic structural stabilization could also contribute to the inhibition of the fibril formation via increasing the apparent activation energy required for the amyloid fibril formation. These two kinetic and thermodynamic effects would realistically work as the factors to suppress the amyloid fibril formation (“*Kinetic and thermodynamic inhibition*” in Figure 8).

## Conclusion

In this study, using TEM and SAXS as well as solid-state NMR spectroscopy and SEC, the detail of the structural development of the prefibrillar intermediates of B chain was investigated. The prefibrillar intermediates showed noodle-like morphology with low β-sheet content, and grew their size and stability through the first and the second prefibrillar intermediates. A new finding was that the second prefibrillar intermediates further elongate, where nucleation tends to be occurring. Furthermore, it was suggested that Fg regulates the length of the prefibrillar intermediates by forming specific complexes with the prefibrillar intermediates. The capability of Fg to form a stable complex with prefibrillar intermediates contributes to suppress the amyloid fibril formation in kinetic and thermodynamic manners. This characteristics of Fg might contribute to reduce the toxicity of various toxic amyloid precursors. Inhibition of the amyloid fibril formation via an interaction with oligomers or prefibrillar intermediates are also suggested among chaperone proteins such as heat shock proteins ^26–29^ and clusterin ^30–32^ as well as non-chaperone proteins,^33, 34^ and thus stabilization of the prefibrillar species by such inhibitor proteins could be a common strategy to inhibit undesirable amyloid fibril formation related to diseases. The detailed effects of Fg on the B chain fibril formation clarified in this study will provide mechanistic insights for the future use of Fg as a potent toxicity neutralizer as well as potent fibril formation inhibitor against amyloid-prone proteins in *in-vivo* systems.

## Materials and Methods

### Purification of B chain and Fg

B chain and Fg were purified in the same manner as described in our previous work with a minor modification.^13^ The detail is describe in Supporting Information (SI).

### Prefibrillar intermediate and amyloid fibril formation

The reaction of fibrillation was initiated by pH jump from ~11 (in 10 mM NaOH) to 8.7 (50 mM Tris-HCl and 5 mM NaCl) at B chain concentration of 1.4 or 2.8 mg/ml. This buffer composition was universally used in this study. In the agitated condition, the sample was shaken at 1,200 rpm at 25 °C using ThermoMixer C (Eppendorf, Germany) so that the fibril formation would be accelerated, whereas no shake was applied in the quiescent condition. When Fg was co-incubated with B chain, the concentration used was 3.5 mg/ml Fg for 1.4 mg/ml B chain, or 7.0 mg/ml Fg for 2.8 mg/ml B chain.

### ThT assay

5 μl sample was mixed with 1ml ThT solution (5 μM ThT, 50 mM Gly-NaOH, pH 8.5), and the fluorescent intensity at 485 nm was monitored with excitation at 445 nm using FP-6500 (JASCO, Japan).

### TEM

A standard negative staining method using uranyl acetate was used. A carbon-supported cupper grid (U1015, EM Japan, Japan) was used for the sample preparation. Briefly, the surface of the grid was loaded on 10 μl sample drop for 1 min, and the drop was removed using a paper. Then the same surface was loaded on 10 μl of uranyl acetate solution (1 % w/w) for 1 min and the solution was removed using a paper. TEM images were taken using HT7700 (HITACHI, Japan). Image analyses were performed by a software, Click Measure (https://onochi-lab.com/sdm_downloads/sdm_downloads-867/). The statistical analyses were conducted using JMP 16.1.0 (SAS Institute Japan).

### SAXS

The SAXS profiles were obtained by using NANOPIX (Rigaku Corporation, Japan). The wavelength of X-ray and the sample-to-detector distance were 0.1542 nm and 1.33 m, respectively. The temperature was kept at 25 °C with a Peltier controller. A transient scattering profile was collected for 30 min and integrated. The scattering vector, *q*, is represented as follows;

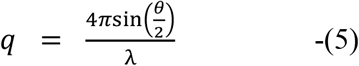

where *λ* and *θ* indicate the X-ray wavelength and the scattering angle, respectively. Under the present set-up, the covered *q* range was from 0.05 to 2.3 nm^-1^. Data acquisition started at 5 min after the reaction was initiated which was continued until the end of the experiments. Data for 30 min were averaged and used for the analyses. The acquisition point was set to be the middle of the time range, and thus the first acquisition time was set to be 20 min. Further information is described in SI.

### Evaluation of structural properties from SAXS data

The intermediate region of the *I*(*q*), which satisfies the condition of 1/*L* < *q* < 1/*D* nm^-1^ (*L* and *D* represent the length and diameter of a rod-like structure, respectively), was analyzed with the following equation;^35^

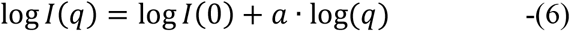

where *I*(0) and *a* represent the intensity at *q* = 0 and the slope of the intermediate region, respectively. The value of the slope (=*a*) was used for evaluating shapes of prefibrillar intermediates. We first fitted at an arbitrary region for obtaining *D*, followed by calculation of *L*. We then checked if these parameters satisfy the condition above. The procedure was repeated until the condition was satisfied.

The cross-section plot for the rod-like structure was analyzed using the equation below; ^35, 36^

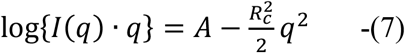

where *A* and *D*_c_ correspond to the constant at *q* = 0 limit and the radius of gyration of the cross section, respectively. The maximum *q* value for the fitting region was determined with the criterion of *R*_c_ *q* < 1.3. We first fitted at an arbitrary region for obtaining *R*_c_, and then checked if the parameters satisfied the requirement. The procedure was repeated until the requirement was satisfied. The base diameter of a rod-like structure, *D*_p_, was obtained as follows;

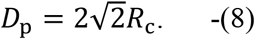

The calculation of the length of prefibrillar intermediates, *L*, was performed based on Broersma’s relationship; ^37^

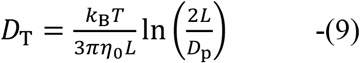

where *D*_T_ is the diffusion coefficient of the solute obtained by DLS (see Supporting Information (SI)). *k*_B_, *T*, and *η*_0_ represent the Boltzmann constant, temperature, and viscosity, respectively. Combing eq. 9 with the Stokes-Einstein-Debye function (SI methods, eq (S1)), we can obtain

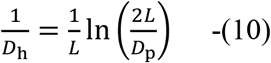

where *D*_h_ indicates the hydrodynamic diameter obtained by DLS. The numerical solution of *L* was obtained using the equation. All the SAXS analyses were performed using Igor Pro 6.37 (Wavemetrics, OR)

### SEC

150 μl sample was analyzed by Superose 6 Inc 10/300 column (Citiva, NY) equipped in ÄKTA purifier (GE healthcare, NY). The elution was conducted at 0.5 ml/min using the same buffer as used for sample at 4 °C

### ^13^C Solid-state NMR spectroscopy

^13^C NMR spectra were recorded on a CMX 400 Infinity NMR spectrometer at the resonance frequencies of 100.0 and 397.8 MHz for carbon and proton nuclei, respectively. CP-MAS and DD-MAS techniques were used to obtain information about secondary structures.^23, 38, 39^ The measurement was performed at 25 °C. More detail is described in SI.

### MALS

SEC-MALS was used to determine the molecular weight of the B chain-Fg complex. A sample was gel-filtrated using the same SEC method above, and elution was further analyzed using a MALS equipment, DAWN 8 (Wyatt Technology, CA) combined with a UV detector, UV-4575 (JASCO, Japan) at 4 °C. The MALS data was analyzed using a software, ASTRA (Wyatt Technology, CA) to calculate the molecular weight.

## Supporting information

Supporting Imformation

## Accession Numbers

human insulin P01308 Fg from bovine plasma P02672 (α chain); P02676 (β chain); P12799 (γ chain)

## Acknowledgements

The SAXS measurement was supported by the Visiting Researchers Program of Institute for Integrated Radiation and Nuclear Science, Kyoto University, and the Project for Construction of the Basis for the Advanced Materials Science and Analytical Study by the Innovative Use of Quantum Beam and Nuclear Sciences at Institute for Integrated Radiation and Nuclear Science, Kyoto University. The TEM images were obtained with the help of Dr. Koki in the department of Histology, School of Medicine, Jichi Medical University. The DLS measurements were supported by Prof. Kuro-o and Prof. Kurosu in Division of Anti-aging Medicine, Center for Molecular Medicine, Jichi Medical University. This research (MALS equipment) was subsidized by JKA through its promotion funds from KEIRIN RACE. MALS measurements were supported by Dr. Unzai (Shoko Science, Japan). This work was conducted under the supports by JSPS Core-to-Core Program, A. Advanced Research Networks. The following JSPS KAKENHI participated in the research: Grant Numbers JP16K17783, JP16H04778, JP16H00772, JP17H06352 and JP20H03224 to E.C., JP17K07361, JP19KK0071, and JP20K06579 to R.I., and JP18H05229, JP18H05534, and JP18H03681 to M.S. N.Y. also thanks to the support by Takeda Science Foundation and Jichi Medical University for the Research Supporting Fund.

## Footnotes

### Supporting Information (SI)

SI methods for purification of B chain, SAXS, DLS, ^13^C solid-state NMR spectroscopy, CD spectroscopy, ^1^H solution NMR spectroscopy, MALS, and calculation of the molecular waeight of a B chain-Fg complex; SI result for DLS; Figure S1, the time-dependent profiles of DLS; Figure S2, (A and B) TEM images of SEC elution fractions of B chain of 1.4 mg/ml at 16 hours and 2.8 mg/ml at 4 hours, respectively; Figure S3, (A-C) Time-course of SAXS profiles of B chain of 2.8 mg/ml and (D) a TEM Image at 4 hours; Figure S4, the time-course of ^1^H NMR measurement at the B chain concentration of 2.8 mg/ml at pH 8.7; Figure S5, (A) curve fitting results of solid-state NMR spectra in the main-chain carbonyl carbon region, and (B) a DD-MAS NMR spectrum in the methyl/methylene region; Figure S6, (A-C) time-course of SAXS profiles of B chain of 1.4 mg/ml incubated with Fg of 3.5 mg/ml, and (D) its SEC result incubated for 18 hours for comparison with Fg only; Figure S7, (A-C) time-course of SAXS profiles of B chain of 2.8 mg/ml incubated with Fg of 7.0 mg/ml, (D) a SEC result B chain 2.8 mg/ml and Fg 7.0 mg/ml, and (E) a TEM image of a SEC elution fraction at ~9 ml indicated in panel D.

### Author Contributions

The manuscript was written through contributions of all authors. All authors have given approval to the final version of the manuscript.

### Notes

The authors declare no competing interests.

