## Supporting Imformation for "Structural development of amyloid precursors in insulin B chain and the inhibition effect by fibrinogen"

Affiliation

### SI methods

**Purification of B chain.** Briefly, human insulin (Wako Pure Chemical Industries, Ltd., Japan) was dissolved in a buffer solution composed of Tris-HCl (56 mM), NaCl (111 mM), and EDTA (2.2 mM), and the pH value was adjusted to 8.6 by HCl. To initiate the reduction step of insulin, 1, 4-dithiothreitol (DTT) solution whose pH value was adjusted to 8.6 by HCl was added to the insulin solution. The final concentration of insulin was 5.0 mg/ml, and the final composition of buffer was 50 mM Tris-HCl, 100 mM NaCl, 2 mM EDTA, and 20 mM DTT. Immediately after the addition of DTT, the solution became turbid because of the precipitation of B chain. The solution was kept under 25 °C for 15 h to complete the precipitation formation. The precipitate was then separated by centrifuging at 17,000 ×*g* for 10 min at 4 °C. The precipitate was then rinsed with Milli-Q® water at 4 °C (Merck Millipore Corp., Germany), centrifuged at 100,000 ×g, and the precipitation was collected. This washing process was repeated three times. The precipitate was dissolved in 10 mM NaOH to give a final concentration of 5.4-6.3 mg/ml, frozen using liquid nitrogen, and stored at -80 °C. The concentration of B chain was determined by using the absorption coefficient of 0.90 $(mg/ml)^{-1}cm^{-1}$ at 280 nm in the NaOH solution. The purity of the B chain was more than 95 %, as checked by the $^{1}H$ signals of ε protons in tyrosine residues obtained using a NMR spectrometer, AVANCEIII HD (Bruker, Germany).

**Purification of Fg.** Fg from bovine plasma (Wako Pure Chemical Industries, Japan) was used for the study. Fg powder of ~50 mg was dissolved in 1ml buffer solution of 100 mM Tris-HCl and 10 mM HCl (pH 8.5), and purified by using a column, HiPrep 16/60 Sephacryl S-300 HR (GE Healthcare, NY) equipped with ÄKTA Start (GE Healthcare,

NY). An elution fraction of Fg monomer was collected and stored at -80 °C.

**SAXS measurement.** To initiate the reactions of the formation of the prefibrillar intermediates, the purified B-chain stock solution in 10 mM NaOH at the concentration of 2.8 mg/ml or 5.6 mg/ml was mixed with the solution of 100 mM Tris-HCl and 10 mM HCl of the same volume so that the final concentration became 1.4 mg/ml or 2.8 mg/ml, respectively. The buffer used in the SAXS experiment was 50 mM Tris-HCl and 5 mM NaCl whose pH value was adjusted to be 8.7. In the case of the presence of Fg, the purified B-chain stock solution in 10 mM NaOH at the concentration of 2.8 mg/ml or 5.6 mg/ml was mixed with Fg/buffer solution of the same volume so that the final concentrations of B chain and Fg became 1.4 mg/ml and 3.5 mg/ml or 2.8 mg/ml and 7.0 mg/ml, respectively. The samples were loaded into a cell with the optical path of ~ 1mm.

**DLS measurement.** In order to estimate particle sizes, DLS experiments were performed using Zetasizer Nano-SZ (Malvern Instruments, Worcestershire, UK). A He-Ne laser at a wavelength of 633 nm was introduced to a sample cuvette and back scattered light was detected using an avalanche photodiode at a scatter angle of 173°. Data collection and processing were performed using the software, Dispersion Technology Software 5.00 (Malvern Instruments Ltd., UK). CONTIN analysis was used for obtaining size distribution based on the Stokes-Einstein-Debye equation;

$$D_{\mathrm{h}} = \frac{k_{\mathrm{B}}T}{3\pi\eta_0 D_{\mathrm{T}}} \quad \text{-(S1)}$$

where $k_{\mathrm{B}}$, $T$, $\eta_0$, and $D_{\mathrm{T}}$ represent the Boltzmann constant, temperature, viscosity, and the experimentally-obtained diffusion coefficient, respectively. The value of the viscosity, $\eta_0$, was set to be that of the buffer solution, i.e. 0.94 mPa·s.

**$^{13}$CSolid-state NMR spectroscopy.** Lyophilized powders were used for the measurement of CP-MAS and DD-MAS NMR spectra. The duration time of the 90 degree pulse was 5.3 μsec, and the repetition time was 4 sec for CP-MAS and DD-MAS experiments. The MAS frequency was set to be 6 kHz. The chemical shift values were calibrated referring to an external carboxyl peak of crystalline glycine at 176.03 ppm from methyl peak of tetramethylsilane (TMS) at 0 ppm. Typically, 5000 transients were acquired and Fourier transformed to obtain CP-MAS and DD-MAS NMR spectra, All the measurements were performed at 25 °C.

**$^{1}$H solution NMR spectroscopy.** NMR spectral measurements were performed using AVANCE III HD equipped with a superconducting magnet that had a Larmor frequency of $^{1}$H of 400.13 MHz (Bruker, Germany). The spectrometer was controlled using the program, Topspin 1.5 (Bruker, Germany). The routine tool Topshim built in Topspin 1.5 was used to maintain high homogeneity of the magnetic field. The spectra were recorded using the zgesgp pulse program that was preinstalled in Topspin 1.5. A 800 μl sample solution was placed in a glass tube with an inner diameter of 4.95 mm, and all experiments were performed at 25 °C. In the fitting of a set of two proton peaks originating from the ε protons of histidine residues in the B chain peptide, a sum of two Lorentzians was used as follows:

$$I(\delta)=\sum_{k=1}^{2}\frac{a_k}{(\delta-\delta_k)^2+\gamma_k} \quad \text{-(S2)}$$

where $I(\delta)$ is peak intensity at the frequency $\delta$, and $a_k$, $\delta_k$, and $\gamma_k$ represent the center amplitude, center frequency, and damping factor of the NMR peak being focused on. The peak area and full width at half-maximum (FWHM) were described as $\pi a_k/\gamma_k$ and $2\gamma_k$, respectively.

**Calculation of the molecular weight of a B chain-Fg complex.** In the case of the complex whose length is ~100 nm, two repeats of seven Fg molecules are expected to exist on the surface of the prefibrillar intermediate. Because the molar ratio of B chain per Fg is ~30, the number of B chain per seven Fg molecules could be ~210. Therefore, the molecular weight can be estimated as follows;

[~340 kDa (Fg) × 7 + ~3.5 kDa (B chain) × 210] × 2 = ~6,200 kDa.

### SI results

#### DLS

The time-course measurement of the hydrodynamic diameter, $D_h$, was performed for the calculation of the length of the rod-like structure. The time-dependent DLS profiles as a function of $D_h$ are shown in Figure S1 at the left side. It is clear that the size is increasing as a function of time. For calculation of the length, the value of $D_h$ at the peak top was used. Its time dependence is shown in figure S1 at the right side. The time dependence was analyzed using model functions as described below.

**B chain 1.4 mg/ml:** At 1.4 mg/ml, $D_h$ increased in a biphasic manner as shown in figure 1B, which we then fit using a bi-exponential function as follows;

$$D_{\mathrm{h}}(\mathrm{t}) = D_{\mathrm{h2}} - (D_{\mathrm{h1}} - D_{\mathrm{h0}})\exp\left(-\frac{t}{\tau_1}\right) - (D_{\mathrm{h2}} - D_{\mathrm{h1}})\exp\left(-\frac{t}{\tau_2}\right) \quad \text{-(S3)}$$

where $D_{h0}$, $D_{h1}$ and $D_{h2}$ represent the hydrodynamic diameter of the initial, first, and second states where the developments from the initial to first state and from the first to second state are characterized by the two time constants of $\tau_1$ and $\tau_2$, respectively. The resultant values were $D_{h0} = 20 \pm 2$ nm, $D_{h1} = 56 \pm 1$ nm, and $D_{h2} = 96 \pm 1$ nm, respectively, with $\tau_1 = 8 \pm 1$ min and $\tau_2 = 460 \pm 30$ min (resultant value ± SD calculated by curve fitting). These values are consistent with those obtained in our previous study where the fast and slow time constants were assigned to the formation of the first and second prefibrillar intermediates, respectively [9].

**B chain 2.8 mg/ml:** At 2.8 mg/ml, $D_h$ increased in a biphasic manner; it steeply increased up to ~200 min, and then gradual increase occurred, which seemed to be almost linear. The time constant of the initial steep increase was then analyzed using a single exponential function;

$$D_{\mathrm{h}}(\mathrm{t}) = D_{\mathrm{h1}} - (D_{\mathrm{h1}} - D_{\mathrm{h0}})\exp\left(-\frac{t}{\tau_1}\right). \quad \text{-(S4)}$$

The fitting was tried in a various fitting range, and we found that up to 200 min the time course was fairly described by the single exponential function. The resultant values were $D_{h0}$ = 24 ± 1 nm, $D_{h1}$ = 93 ± 3 nm, respectively, with $\tau_1$ = 88 ± 9 min (resultant value ± SD calculated by curve fitting). The value of $D_{h1}$ is close to $D_{h2}$ obtained in the analysis of 1.4 mg/ml, which was assigned as the hydrodynamic diameter of the second prefibrillar intermediate. This fact supports the idea that the initial event at 2.8 mg/ml corresponds to the formation of the second prefibrillar intermediate. The Further linear increase in the hydrodynamic diameter thus indicates a further structural development from the second prefibrillar intermediate, which was not observed at 1.4 mg/ml.

**B chain 1.4 mg/ml+Fg 3.5 mg/ml:** The value of $D_h$ increased in a biphasic manner; the value steeply increased up to 30 min followed by a further slow increase. The time-dependence was then fitted using eq S3, resulting in obtaining $D_{h0}$ = 22 ± 1 nm, $D_{h1}$ = 33 ± 1 nm, and $D_{h2}$ = 42 ± 1 nm, respectively, with $\tau_1$ =15 ± 3 min and $\tau_2$ = 370 ± 50 min (resultant value ± SD calculated by curve fitting). In our previous study, qualitatively similar time constant to $\tau_1$ was reported in CD spectroscopy (21 ± 3 min) [11]. Therefore, the first phase is assumed to be the formation of the first prefibrillar intermediate. On the contrary, the value of $\tau_2$ is much shorter than that obtained using CD spectroscopy in our previous study. This discrepancy might be because the structural development of the second prefibrillar intermediate is decoupled from its further secondary structure formation for the development of the second prefibrillar intermediate. More detail is discussed in the main text.

**B chain 2.8 mg/ml+Fg 7.0 mg/ml:** $D_h$ increased in a biphasic manner; a slight increase was observed up to ~100 min, follow by a further slow increase. The time course was then fitted using the biexponential function as represented by eq. S3. As a result, the resultant values were $D_{h0} = 29 \pm 1$ nm, $D_{h1} = 33 \pm 1$ nm, and $D_{h2} = 41 \pm 1$ nm, respectively, with $\tau_1 = 92 \pm 27$ min and $\tau_2 = 1{,}040 \pm 420$ min (resultant value ± SD calculated by curve fitting). The value of $D_{h1}$ and $D_{h2}$ are almost the same those obtained in the case of B chain 1.4 mg/ml and Fg 3.5 mg/ml (Fig. 4). This fact, together with the fact that the base radius were almost identical in these two different concentration conditions, strongly indicates that the same first and second prefibrillar intermediates were formed even in these two different concentrations.

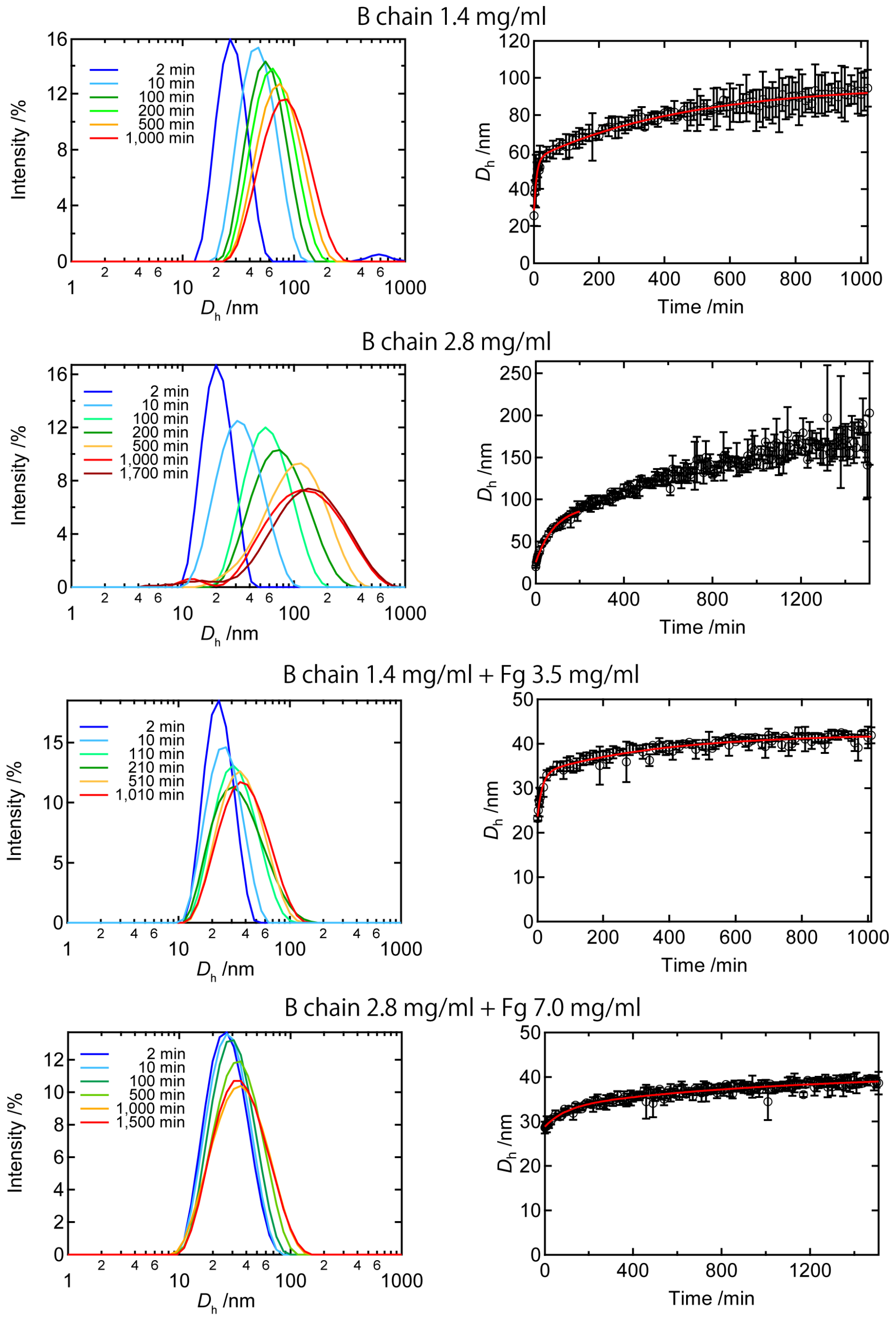


**Figure S1**. Time-dependent DLS profile as a function of $D_h$ (left) and time-dependence of $D_h$ at the peak of the profile (right). In the right panels, the red line shows simulated curves obtained by fitting using several functions as described above. We used the value of $D_h$ at the peak of the DLS intensity profile for the analysis. Hereafter, $D_h$ indicates the peak of the DLS intensity profile unless otherwise noted.

A

B

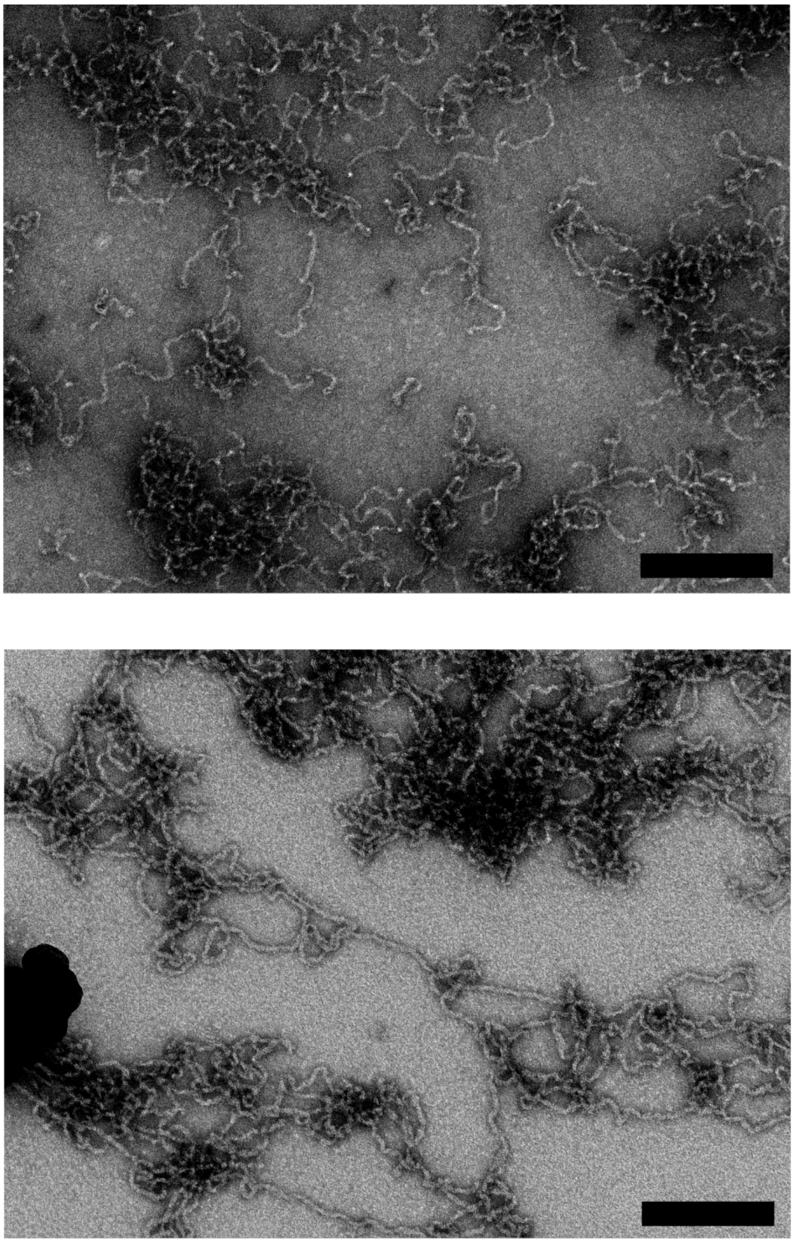

**Figure S2**. TEM images of B chain of 1.4 mg/ml at (A) 30 min and (B) 16 hours, respectively. The scale bars indicate 200 nm.

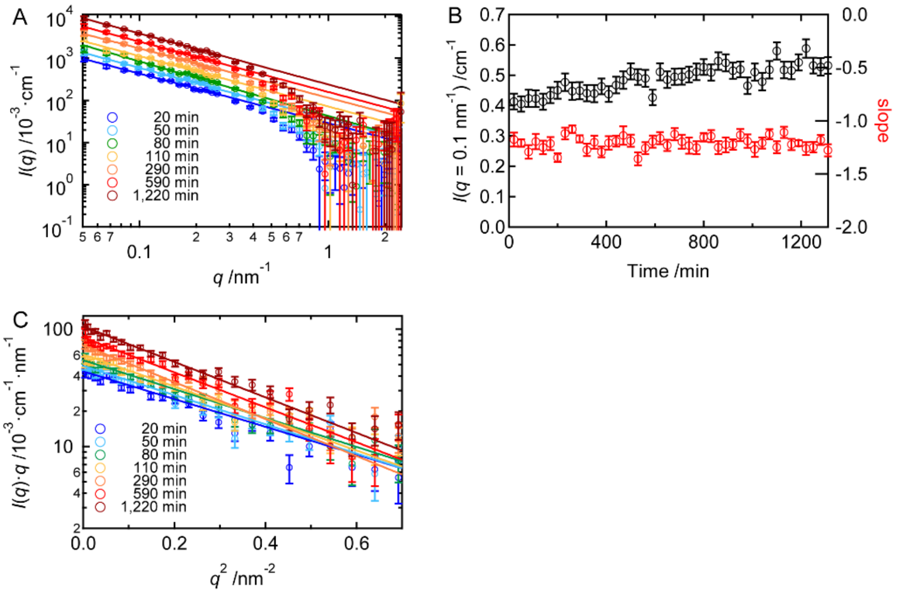


**Figure S3**. (A-C) Time-course of SAXS profiles of B chain of 2.8 mg/ml. (A) The intensity vs $q$ plot. The lines represent fitting results in the intermediate region obtained by using eq. 6 at 0.07 $\leq q \leq$ 0.1 nm$^{-1}$. Each plot is arbitrary shifted for guiding eyes. (B) Time-dependence of the intensity at $q$ = 0.1 nm$^{-1}$ (black symbol) and the slope of the fitting lines shown in A, respectively. The initial intensity was already high compared to the case of 1.4 mg/ml, and moderately increased as a function of time. This tendency is different from that of the case of 1.4 mg/ml where the drastic increase from the low initial intensity was observed (Figure 3A and B), representing that the formation of the prefibrillar intermediates was indeed accelerated upon increasing B chain concentration. (C) The cross-section plot. The lines represent fitting results performed using eq. 7 at 0.12 $\leq q \leq$ 0.16 nm$^{-1}$ to obtain the base radius of gyration. Each plot is arbitrary shifted for guiding eyes. See detail in the main text.

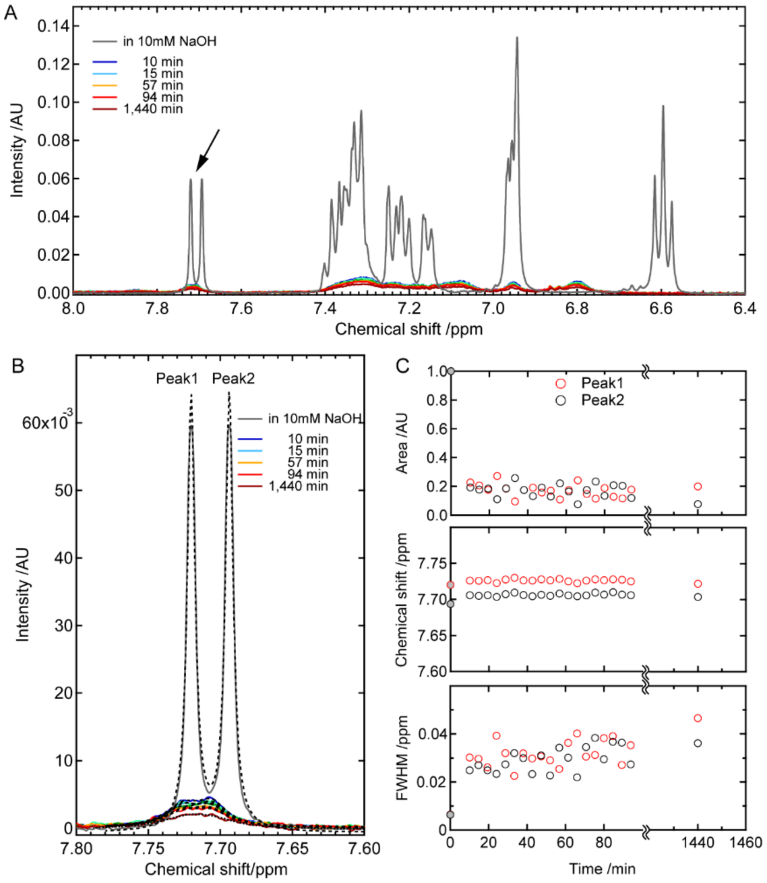


**Figure S4**. The time-course of $^{1}$H NMR measurement at the B chain concentration of 2.8 mg/ml at pH 8.7. (A) Time-dependent spectra in the low-magnetic-field region. The spectrum in 10 mM NaOH condition where monomeric B chain only exists is also shown. The arrow indicates histidine $\varepsilon$ proton peaks of two different histidine residues. (B) A magnified view of the histidine $\varepsilon$ proton peaks. These two peaks are designated as peak1 and peak2, which are marked above each peak. These two peaks were fitted by using eq. S2, and the resultant simulated waves are shown by the black dotted lines. (C) Areas, chemical shifts, and FWHMs of the histidine $\varepsilon$ peaks obtained from the curve fitting. The values of 10 mM NaOH condition, which are colored by gray, are shown at time zero. The monomeric component quickly decreased to ~20 % within 10 min after the initiation of reaction and remained constant. This result supports the idea that the formation of the first prefibrillar intermediate instantaneously completed within the experimental dead time of the time-resolved SAXS measurement. The fraction of the prefibrillar intermediates, ~80 %, was higher than that of the case of 1.4 mg/ml (~60 %, obtained in our previous study), representing that the increase in the B chain concentration also contributed to increase the amount of the prefibrillar intermediates.

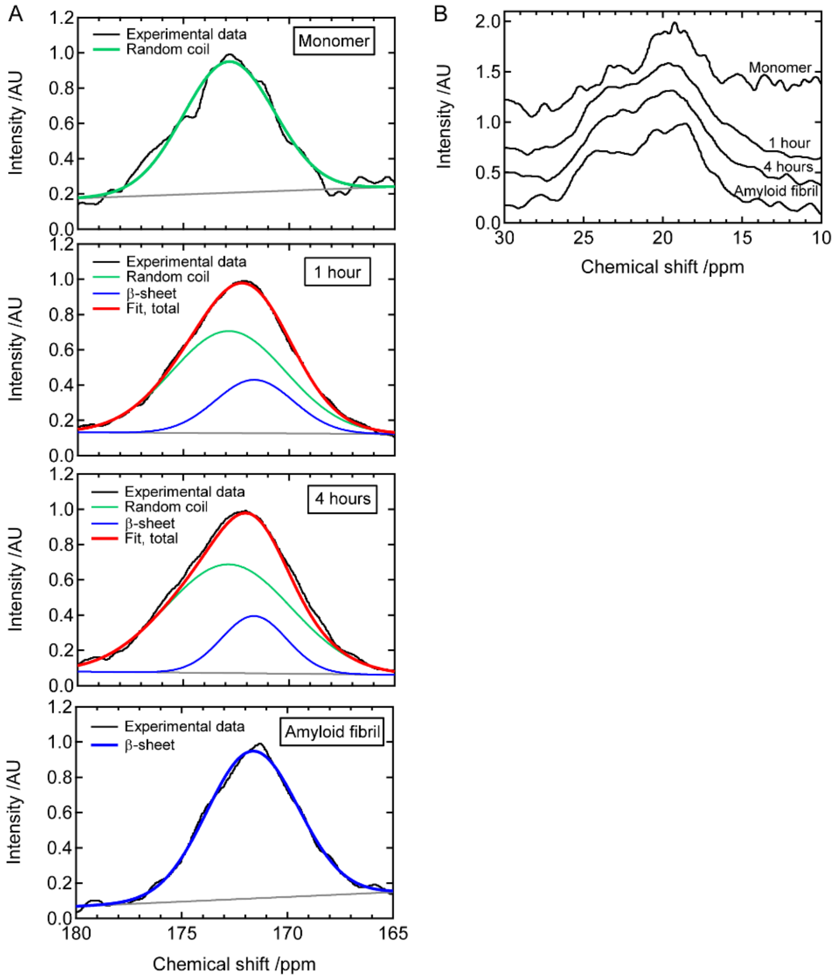


**Figure S5**. (A) The fitting results of the solid-state NMR spectra to estimate secondary structural components. The spectra of the prefibrillar intermediates were fitted by a linear combination of two Gaussian functions where the first peak position was fixed to that of the random-coil state and second peak position to that of the amyloid-fibril structure. Under these restrictions, a global fitting was performed on these four spectra. (B) The DD-MAS NMR spectrum in the methyl/methylene region. See detail in the main text.

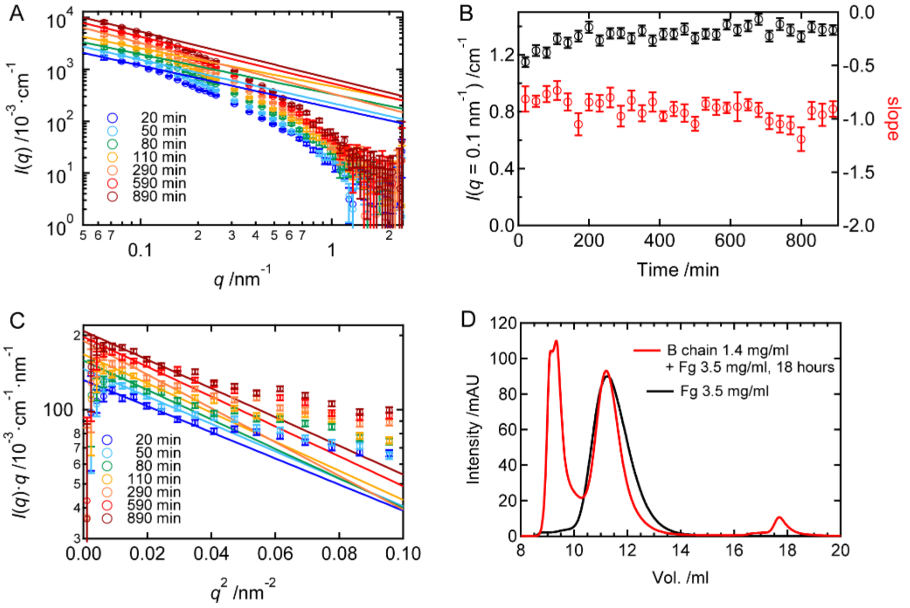


**Figure S6**. (A-C) Time-course of SAXS profiles of B chain of 1.4 mg/ml incubated with Fg of 3.5 mg/ml. (A) The intensity vs $q$ plot. The lines represent fitting results obtained by using eq. 6 at $0.07 \leq q \leq 0.1$ nm$^{-1}$. Each plot is arbitrary shifted for guiding eyes. (B) Time-dependence of the intensity at $q = 0.1$ nm$^{-1}$ (black symbol) and the slope of the fitting lines shown in A, respectively. The scattering intensity gradually increased as a function of time. The slope was at around -1 constantly, representing formation of rod-like structures. (C) The cross-section plot. The lines represent fitting results performed using eq. 7 at $0.1 \leq q \leq 0.156$ nm$^{-1}$ to obtain the base radius of gyration. Each plot is arbitrary shifted for guiding eyes. See detail in the main text. (D) Comparison of SEC results at B chain 1.4 mg/ml incubated with Fg 3.5 mg/ml for 18 hours and Fg only.

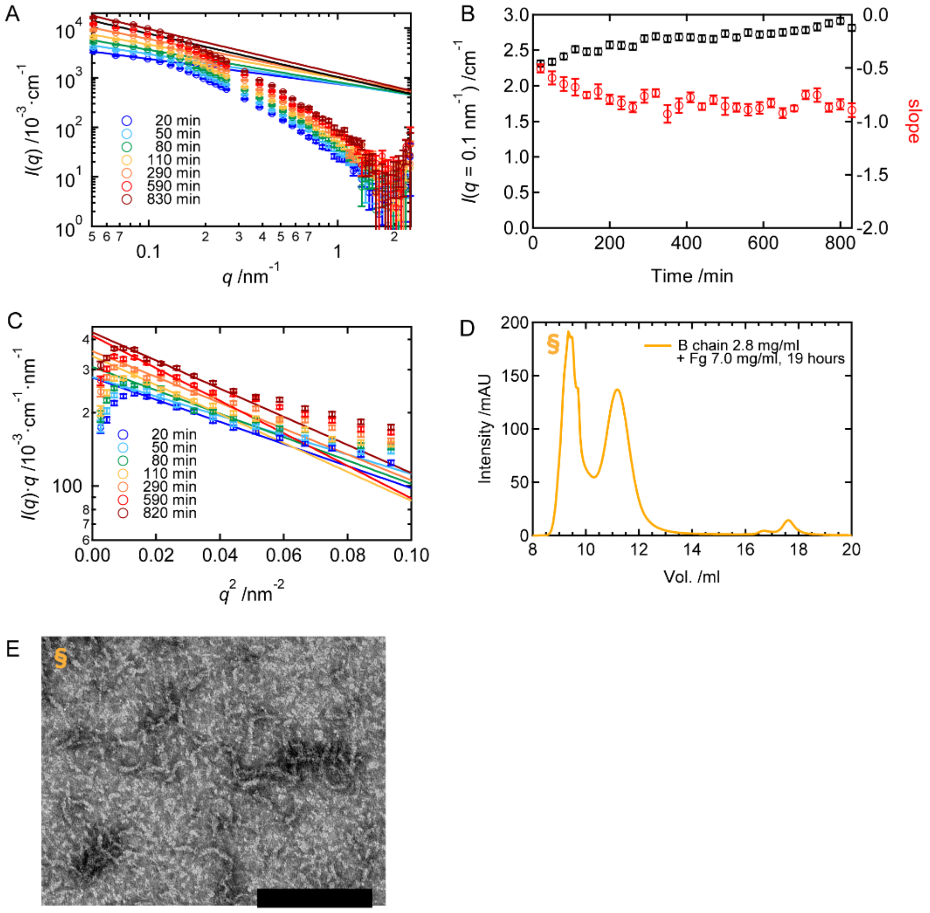


**Figure S7**. (A-C) Time-course of SAXS profiles of B chain of 2.8 mg/ml incubated with Fg of 7.0 mg/ml. (A) The intensity vs *q* plot. The lines represent fitting results obtained by using eq. 6 at $0.07 \le q \le 0.1$ nm$^{-1}$. Each plot is arbitrary shifted for guiding eyes. (B) Time-dependence of the intensity at $q = 0.1$ nm$^{-1}$ (black symbol) and the slope of the fitting lines shown in A, respectively. (C) The cross-section plot. The lines represent fitting results performed using eq. 7 at $0.12 \le q \le 0.16$ nm$^{-1}$ to obtain the base radius of gyration. Each plot is arbitrary shifted for guiding eyes. See detail in the main text. (D) A SEC result of B 2.8 mg/ml B chain incubated with 7.0 mg/ml Fg incubated for 19 hours. (E) A TEM image of ~9 ml elution in the SEC result as indicated by § in panel D.
